# Structure and function of the Si3 insertion integrated into the trigger loop/helix of cyanobacterial RNA polymerase

**DOI:** 10.1101/2024.01.11.575193

**Authors:** M. Zuhaib Qayyum, Masahiko Imashimizu, Miron Leanca, Rishi K. Vishwakarma, Amber Riaz-Bradley, Yulia Yuzenkova, Katsuhiko S. Murakami

## Abstract

Cyanobacteria and evolutionarily related chloroplasts of algae and plants possess unique RNA polymerases (RNAPs) with characteristics that distinguish from canonical bacterial RNAPs. The largest subunit of cyanobacterial RNAP (cyRNAP) is divided into two polypeptides, β’1 and β’2, and contains the largest known lineage-specific insertion domain, Si3, located in the middle of the trigger loop and spans approximately half of the β’2 subunit. In this study, we present the X-ray crystal structure of Si3 and the cryo-EM structures of the cyRNAP transcription elongation complex plus the NusG factor with and without incoming nucleoside triphosphate (iNTP) bound at the active site. Si3 has a well-ordered and elongated shape that exceeds the length of the main body of cyRNAP, fits into cavities of cyRNAP and shields the binding site of secondary channel-binding proteins such as Gre and DksA. A small transition from the trigger loop to the trigger helix upon iNTP binding at the active site results in a large swing motion of Si3; however, this transition does not affect the catalytic activity of cyRNAP due to its minimal contact with cyRNAP, NusG or DNA. This study provides a structural framework for understanding the evolutionary significance of these features unique to cyRNAP and chloroplast RNAP and may provide insights into the molecular mechanism of transcription in specific environment of photosynthetic organisms.

**Significance statement:** Cellular RNA polymerase (RNAP) carries out RNA synthesis and proofreading reactions utilizing a mobile catalytic domain known as the trigger loop/helix. In cyanobacteria, this essential domain acquired a large Si3 insertion during the course of evolution. Despite its elongated shape and large swinging motion associated with the transition between the trigger loop and helix, Si3 is effectively accommodated within cyRNAP, with no impact on the fundamental functions of the trigger loop. Understanding the significance of Si3 in cyanobacteria and chloroplasts is expected to reveal unique transcription mechanism in photosynthetic organisms.

## Introduction

Cyanobacteria and chloroplasts of algae and higher plants are characterized by oxygen-evolving photosynthesis and are phylogenetically closely related. These genomes are transcribed by a bacterial-type RNA polymerase (cyRNAP and plastid-encoded RNAP, PEP, respectively) aided by transcription initiation σ factors for recognition of specific promoters (1–3). Although cyRNAPs and chloroplast PEPs retain the fundamental functions of bacterial RNAPs, they possess several distinct characteristics that distinguish them from canonical bacterial RNAPs.

First, the largest subunit of cyRNAP is separated into two polypeptides, β’1 and β’2, which are encoded by the *rpoC1* and *rpoC2* genes, respectively (Fig. 1A). In *Synechococcus elongatus*, which is the cyanobacterium used for the cryo-EM structural study of RNAP described herein, the 624 residue β’1 and 1,318 residue β’2 subunits correspond to the amino (N)-terminal one-third and the carboxy (C)-terminal two-thirds of the 1,407 residue β’ subunit in *Escherichia coli*, respectively. A junction between the β’1 and β’2 subunits is positioned before the conserved region E (4, 5). The β’1 subunit contains the clamp and the catalytic double-psi-β-barrel domain coordinating a Mg^2+^ ion; the β’2 subunit contains the rim helix, bridge helix, trigger loop and jaw domain.

**Fig. 1.**
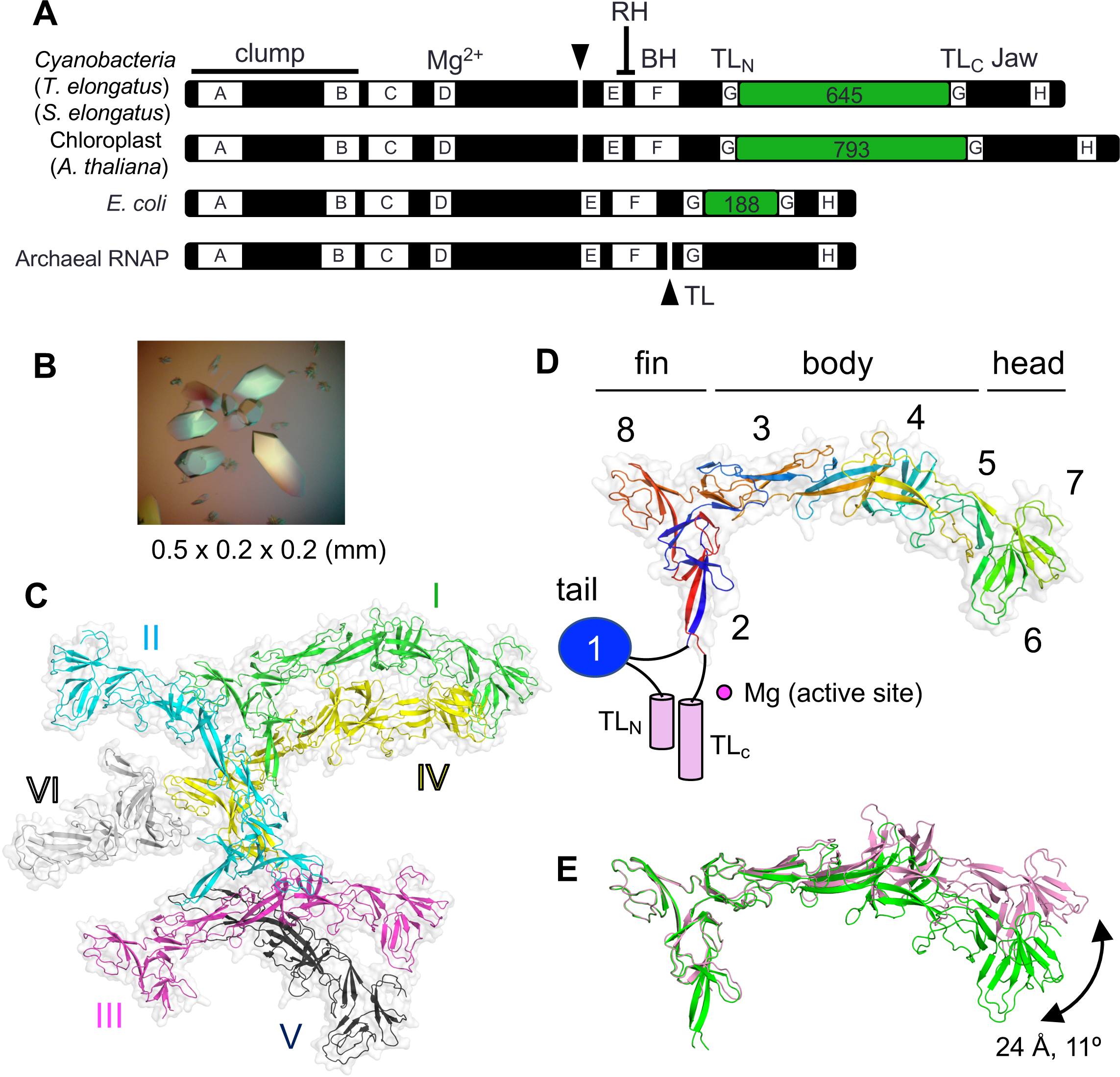
X-ray crystal structure of TelSi3ΔN. (A) The thick bars represent the primary sequences of the largest subunits of the bacterial, chloroplast and archaeal RNAPs. Domains (Si3, green boxes) and structural motifs (RH, rim helix; BH, bridge helix; TL, trigger loop) are labeled. The lettered boxes represent evolutionarily conserved regions. The split ends of the two polypeptides are indicated by black triangles. (B) Crystals of TelSi3ΔN. (C) Structure of TelSi3ΔN. Six molecules of TelSi3ΔN (I∼VI) are present in the asymmetric unit. Molecules are depicted as cartoon models with transparent surfaces, and each molecule is denoted by a unique color and labeled. (D) The backbone is colored as a ramp from the N-terminus to the C-terminus, from blue/cyan/green/yellow/orange/red. SBHMs are labeled 1 to 8, and subdomains (tail, fin, body and head) are indicated. The TelSi3ΔN structure lacks SBHM-1, and the trigger loops (TL_N_ and TL_C_) are depicted as blue oval and pink cylinders, respectively, with black lines showing their connections with TelSi3ΔN. (E) Molecules 1 and 3 of TelSi3ΔN are superimposed via fin subdomains, revealing flexibility in the orientation between the fin and body/head subdomains.

Second, cyRNAP contains the largest known lineage-specific insertion domain, Si3 (645 residues), which spans approximately half the size of the β2’ subunit and is located in the middle of the trigger loop (Fig. 1A) (6, 7). The trigger loop plays a central role in nucleotide selection, RNA synthesis and RNA cleavage during proofreading by cellular RNAPs (8). In the absence of nucleotide triphosphate (NTP) substrate, the tip of the trigger loop is located away from the active site (9). Upon binding of complementary incoming NTP (iNTP) at the active site, the trigger loop folds to form a trigger helix containing two α-helices, which extensively interacts with the base and triphosphate groups of iNTP and facilitates the nucleotidyl transfer reaction (8). The Si3 insertion is found in RNAPs of gram-negative bacteria in the middle of the trigger loop (evolutionarily conserved region G; Fig. 1A). Si3 is composed of repeats of the conserved sandwich-barrel hybrid motif (SBHM). *Escherichia coli* (*E. coli*) RNAP contains two copies of SBHM (SFig. 1A) (10), and sequence analysis indicates that up to seven copies of SBHM are present in the Si3 insertion of cyRNAP (6). The structure and function of the Si3 insertion in *E. coli* RNAP have been well characterized; it is involved in stabilizing the open complex and RNA hairpin-dependent (*his*) and -independent (*ops*) transcription pausing (11, 12) and is highly mobile, with its confirmation being dependent on the folded/unfolded state of the trigger loop/helix and binding of transcription factors (Gre, DksA) at the secondary channel of RNAP (13, 14). In addition, structural and functional analyses of Si3 in cyRNAP have recently been initiated. According to the cryo-electron microscopy (cryo-EM) structure of the cyRNAP promoter complex (15), Si3 forms an “arch” with region 2 of the σ factor, the element involved in opening the DNA duplex at the -10 position of the promoter. This arch stabilizes the promoter complex, and its removal affects the fitness and stress resistance of cyanobacteria. Notably, the Si3-σ contact remains intact upon trigger loop refolding into the trigger helix after iNTP addition to the initiation complex with the short RNA transcript. After transition to the elongation phase, it is unknown whether Si3 becomes mobile in the presence of transcription elongation factors such as NusG and how Si3 affects refolding of the trigger helix and the catalytic activity of RNAP.

In this work, we structurally and biochemically analyzed cyRNAP elongation complex (EC) to understand the functional importance of Si3 in the elongation phase of transcription. We solved the X-ray crystal structure of Si3 and cryo-EM structures of the cyRNAP EC with NusG in the presence and absence of iNTP bound at the active site.

## Results

### X-ray crystal structure of *Thermosynechococcus elongatus BP-1* Si3 (TelSi3)

We investigated the structure of the separate Si3 protein of the thermophilic cyanobacterium *Thermosynechococcus elongatus BP-1* (TelSi3) by X-ray crystallography. The DNA sequence encoding Si3 (residues 345-983) was cloned and inserted into a vector for expression in *E. coli* cells, and the resulting protein was purified to homogeneity. Initial attempts to crystallize TelSi3 were unsuccessful. Limited trypsinolysis revealed that the amino-terminal (N-terminal) 91 residues of TelSi3 are sensitive to proteolysis (SFig. 2A), indicating flexibility, which potentially hindered crystallization. We then cloned and expressed TelSi3, which lacks the N-terminal 91 residues (TelSi3ΔN, residues 435 to 983) and thus forms large crystals (Fig. 1B) belonging to the P3(2)21 space group (six TelSi3ΔN copies per asymmetric unit; Fig. 1C). We were unable to generate a TelSi3ΔN model suitable for molecular replacement based on the protein sequence (e.g., by SWISS-MODEL; SFig. 2C). Therefore, the experimental phase was achieved by the single-wavelength anomalous dispersion (SAD) method using selenomethionine (SeMet)-labeled TelSi3ΔN protein (SFig. 2B). The 3.2 Å resolution experimental density map allowed us to build the structures of four full-length and two partial models of TelSi3ΔN in the asymmetric unit (STable 1). The AlphaFold (20) structural prediction for TelSi3ΔN was in close agreement with the X-ray structure, with an RMSD of 1.08 Å (SFig. 2C).

TelSi3ΔN (150 Å in length and 50 Å in width) is longer than the canonical bacterial RNAP (e.g., 110 × 130 Å: *E. coli* RNAP) (SFig. 1B). TelSi3ΔN comprises seven SBHMs (SBHM-2 to SBHM-8). The X-ray crystal structure of the N-terminal region (81 residues) of Si3 from *S. elongatus* PCC 7942 (15) showed an independently folded SBHM (SBHM-1). This region corresponds to the 91 N-terminal residues of TelSi3 (missing in the crystallized TelSi3ΔN), indicating that cyRNAP contains 8 copies of SBHM within Si3 (Fig. 1C, SFig. 3).

TelSi3 has a swordfish-shaped profile, with distinct “tail”, “fin”, “body” and “head” subdomains formed by SBHM-1, SBHM-2/8, SBHM-3/4/5 and SBHM-6/7, respectively (Fig. 1D). Notably, the SBHMs in TelSi3 are not structured in a simple tandem arrangement (Fig. 2D and SFig. 3), in contrast to *E. coli* Si3, which contains two independently folded SBHMs connected by a short linker (SFig. 1A) (10). Although each SBHM has a core antiparallel β-sheet topology, connections between the β-sheets vary as the polypeptide chain folds over itself (Fig. 1D). In addition, the sequences of SBHM-1, -6, -7 and -8 are continuous; the others (SBHM-2, - 3, -4 and -5) contain structural elements from distant regions of the polypeptide sequence. We assessed conformational flexibility by comparing the four full-length TelSi3ΔN structures from the asymmetric unit using the Si3-fin as a reference for superimposition. This showed substantial conformational variation in the Si3-head, allowing for a 24 AL displacement associated with an 11° rotation (Fig. 1E).

**Fig. 2.**
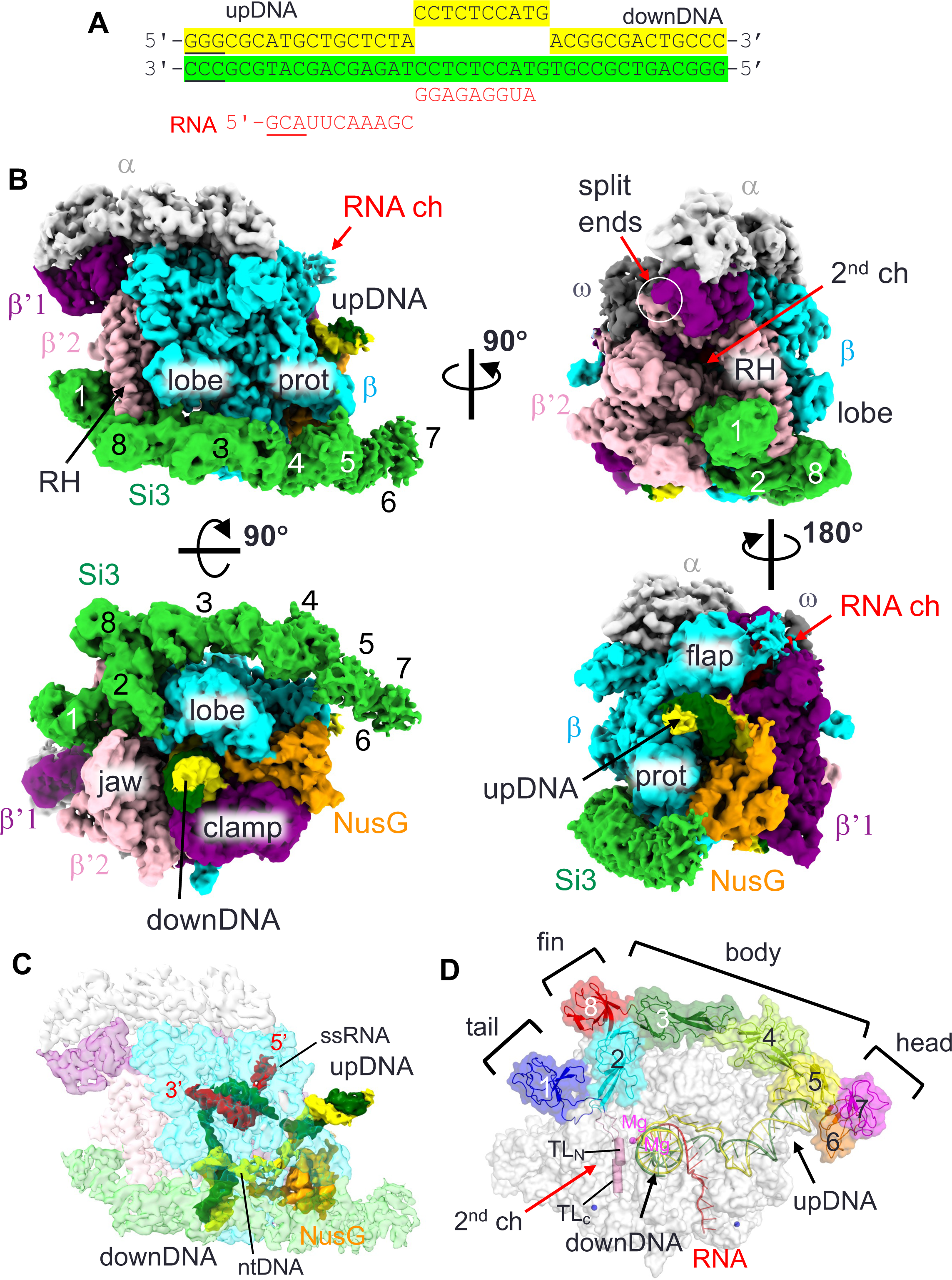
Cryo-EM image of the cyRNAP elongation complex with NusG. (A) The sequence of the DNA/RNA scaffold used for the EC-NusG assembly (template DNA, green; nontemplate DNA, yellow; RNA, red). DNA and RNA regions lacking cryo-EM density are underlined. (B) Orthogonal views of the cryo-EM density map. Subunits and domains of cyRNAP, DNA, RNA and NusG are colored and labeled (RH, rim helix; prot, protrusion; downDNA, downstream DNA; upDNA, upstream DNA). The split ends of the β’1 and β’2 subunits are indicated by white circles. The SBHMs in Si3 are labeled 1 to 8. (C) Cryo-EM density of DNA, RNA and NusG are shown with a transparent RNAP density map (ntDNA, nontemplate DNA; ssRNA, single-stranded RNA). The 5’ and 3’ ends of the RNA are indicated. The cryo-EM density map is colored according to B. (D) Efficient storage of an elongated and large Si3 molecule on the surface of cyRNAP. The structure of EC-NusG is shown as a transparent surface, and the Si3, DNA/RNA and trigger loop (TL_N_, TL_C_) regions are shown as cartoon models. SBHMs are labeled 1 to 8, and subdomains (tail, fin, body and head) are indicated. The active site of RNAP is designated by catalytic Mg^2+^ (magenta sphere).

### Cryo-EM structure of the *Synechococcus elongatus* RNAP elongation complex with NusG

To investigate the structure of cyRNAP and the dynamics of Si3 at the elongation stage, we determined the cryo-EM single-particle reconstruction structure of the cyRNAP EC (SFig 4, STable 2). NusG was also included in the EC, as most ECs contain NusG under physiological transcription conditions (16, 17), and physical contact between Si3 and NusG was investigated.

We used recombinant double affinity-tagged *S. elongatus* cyRNAP to avoid isolation of any chimeric cyRNAP containing *E. coli* RNAP subunit. EC was assembled by mixing cyRNAP, NusG and the DNA/RNA scaffold (Fig. 2A). The preferred particle orientation issue of EC-NusG was resolved by adding CHAPSO (final concentration of 0.8 mM) to the sample before application to the cryo-EM grid (18). The cryo-EM structure was determined with an overall resolution of 3 AL, revealing well-defined cryo-EM densities for cyRNAP, the N-terminal domain of NusG (residues 19-138) and the DNA/RNA hybrid (Fig. 2B). The densities of the single-stranded nontemplate DNA in the transcription bubble and the single-stranded RNA within the RNA exit channel were traceable due to their respective interactions with NusG and the RNA exit channel (Fig. 2C). The carboxyl-terminal (C-terminal) domains of the α subunits and the Kyrpides-Ouzounis-Woese (KOW) domain of NusG were disordered.

By contacting both the upstream and downstream DNA duplexes, NusG seemed to maintain a 90° bend in the DNA centered at the RNAP active site (SFig. 5A), which may stabilize the DNA/RNA holding of cyRNAP. To evaluate the role of NusG, we immobilized reconstituted ECs on agarose beads and challenged the complex with 300 mM NaCl in the absence of NusG. There was a significant reduction in the proportion of RNA released from the complex compared with the EC in the presence of NusG (SFig. 5B), indicating its stabilizing effect. Notably, compared with its orthologs from *E. coli*, *Bacillus subtilis* and *Mycobacterium tuberculosis*, the cyanobacterial NusG gene possesses a longer and more positively charged loop (residues 110-122) within the N-terminal domain. This loop extends toward the downstream DNA and single-stranded non-template DNA within the transcription bubble (SFig. 5A). Deletion of this cyanobacteria-specific loop (NusG^Δ110-122^) significantly reduced the stabilizing effect of NusG (SFig. 5B).

### Si3 runs along the cavities of cyRNAP and shields the binding site of DksA/Gre factors

By fitting the models of RNAP (without Si3), NusG and the DNA/RNA scaffold, we elucidated a density corresponding to Si3, which extends starting from the trigger loop and then moves below the rim helix (β’2 subunit), running along the lobe/protrusion domains (β subunit) and nearly reaching the upstream DNA (Fig. 2B, SMovie 1). The overall structure of cyRNAP is nearly identical to the structures of other bacterial RNAPs, including those of *E. coli* and *M. tuberculosis* (19, 20), indicating that Si3 runs along the cavities of RNAP without influencing its general shape or conformation. The crystal structures of Si3 containing both SBHM2-8 and SBHM1 were fitted to their corresponding cryo-EM density. The cryo-EM density of the Si3-head was weak and had a low resolution (Fig. 2B, SFig. 4E), suggesting its mobility.

Si3-tail is positioned in front of the rim helix (Fig. 3A). Si3-fin is positioned below the rim helix, and the extended SBHM2 loop (residues 463-471) fills a gap between the β’2 jaw and β lobe domains. Si3-body is located beside the lobe and protrusion domains of the β subunit, and Si3-head reaches the upstream DNA (Figs. 2B and 3A). Si3-fin contacts the bottom part of the rim helix, but only a few amino acid residues of Si3 contact the main body of RNAP and NusG, suggesting that Si3-tail and Si3-body/head can move their positions without restraint. Si3 spans the entire length of cyRNAP, reaching from the secondary channel to the upstream DNA. However, it likely does not interfere with any basic function of cyRNAP (i.e., DNA binding, RNA elongation, binding of initiation factor σ, or elongation factors NusA and NusG), as it runs along the sidewall of cyRNAP (Figs. 2D and 3A).

**Fig. 3.**
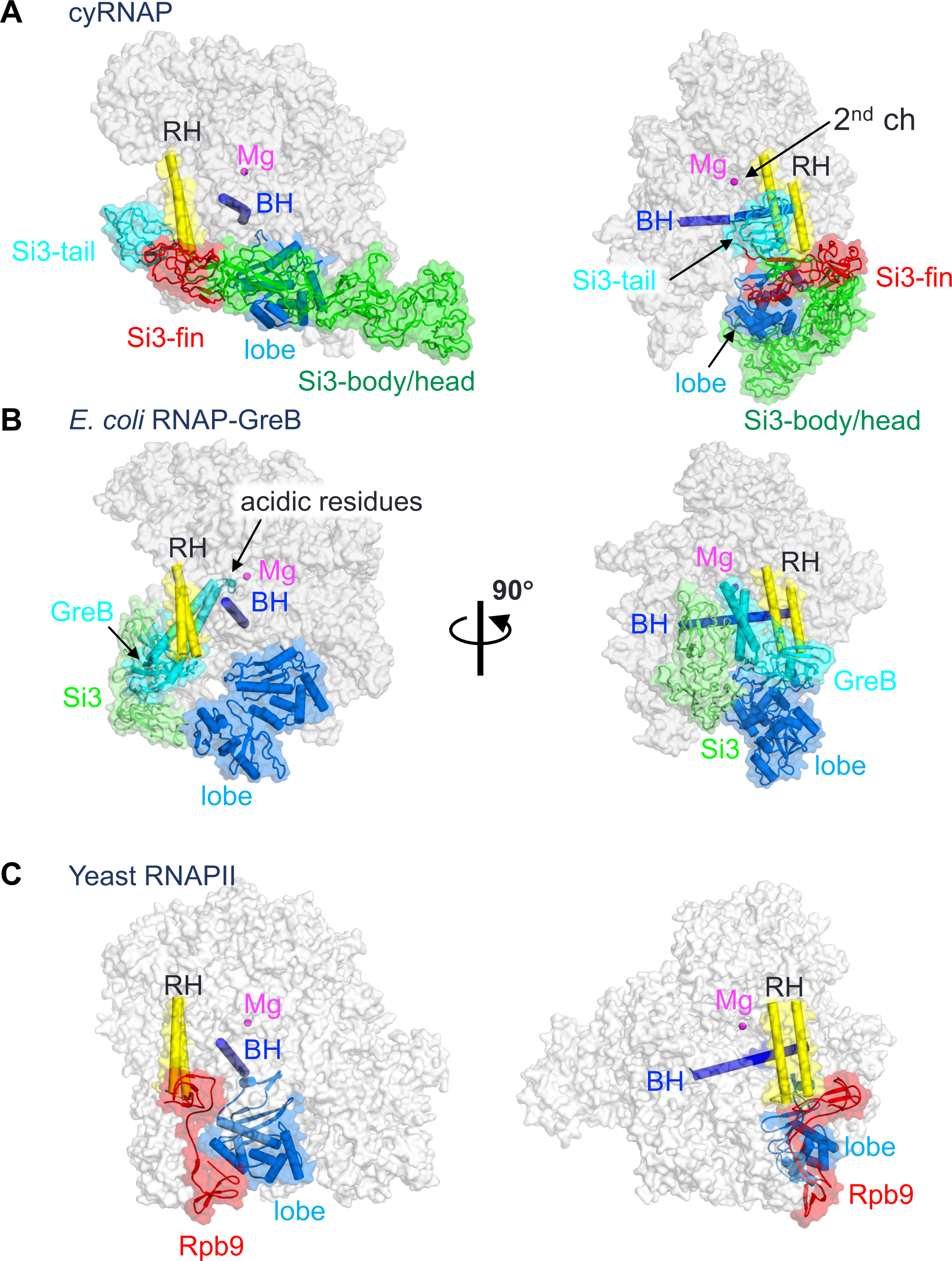
Comparison of the structures of cyRNAP, *E. coli* RNAP and eukaryotic RNAPII. The structures of cyRNAP (**A**), *E. coli* RNAP with GreB (PDB: 6RIN, **B**) and yeast RNAPII (PDB: 7ML0, **C**) are shown as transparent surfaces with domains, subunits and a factor described in the main text.

During transcription, the secondary channel of all cellular RNAPs, including bacterial RNAPs, serves as the only access route between the active site found in the center of RNAP and the external milieu, serving as an entry point for substrate NTPs and an exit route for the RNA 3’-end during backtracking (prior to RNA cleavage). In cyRNAP, the secondary channel appears to be open enough to allow these functions. In addition to these basic functions, the secondary channel serves as a binding platform for proofreading factors such as Gre and regulatory factors such as DksA, known as secondary channel binding factors (13, 27). These factors use the RNAP rim helix as a primary binding site, after which the coiled-coil domain is inserted to access the active site of RNAP (Fig. 3B). In cyRNAP, Si3-tail and -fin occupy the front and bottom sides of the rim helix, respectively, thereby preventing any potential association of secondary channel binding factors (Fig. 3A).

### Dynamic motion of Si3 associated with the transition between the trigger loop and helix during iNTP binding at the active site

To investigate the Si3 conformational change associated with trigger helix refolding, we prepared an iNTP-bound form of the EC by extending RNA with 3’-deoxy adenosine triphosphate (3’-dATP), which arrested further RNA extension, followed by cytosine triphosphate (CTP) addition as the iNTP (SFig. 6). The resulting cryo-EM structure was determined at 2.79 Å resolution (SFig. 6). Although an excess amount of CTP was added to the EC, a substantial population of ECs (∼40%) remained unbound to iNTP. However, the iNTP-bound EC could be clearly distinguished from the iNTP-free EC during 3D classification of the cryo-EM data process due to its unique Si3 orientation relative to the main body of cyRNAP associated with iNTP binding (SFig. 6B and 6D). This allowed for a well-defined density map of the cyRNAP active site. In the iNTP-bound EC, the B-site Mg^2+^ (known as the nucleotide-binding metal) was present at the active site. However, the A-site Mg^2+^ (known as the catalytic metal) was absent, likely due to the lack of a hydroxyl group at the 3’-end of the RNA. Trigger helix folding establishes several essential contacts between the iNTP and amino acid residues, including β’2-M339 in contact with the nucleobase and β’2-H343 in contact with the β-phosphate group (SFig. 6D).

Trigger helix folding induces significant motion of Si3 relative to the main body of cyRNAP. Specifically, the trigger helix formation pulls a linker connecting the C-terminal half of the trigger helix and the Si3-fin, and during this process, the tip of the rim helix acts as a pivot point, converting the lateral motion of the linker (∼10 Å) into the rotational motion of Si3, resulting in an ∼50 Å distance and a 24° swing of Si3-head (Figs. 4A and B, SMovie 2). Si3-body/head swings down from the main body of cyRNAP; thus, the β protrusion domain no longer contacts Si3-body/head in the iNTP-bound EC (Fig. 4A). Remarkably, the large swinging of Si3, which is coupled to trigger helix formation (Fig. 4B), did not markedly alter the catalytic properties of cyRNAP (Fig. 4C). Three ECs containing 14, 15 and 16 nucleotide long RNAs (EC14, 15 and 16) were prepared by extending the initial 5’-labelled 13 nt long RNA in the nucleic acid scaffold shown above the summary table. Nucleotide addition, its direct reversal by pyrophosphorolysis, and transcript cleavage were performed for the ECs that formed with either wild-type (WT) or Si3-lacking (ΔSi3) cyRNAP. Rates of the NTP addition, pyrophosphorolysis and RNA hydrolysis were similar between the WT and ΔSi3 cyRNAPs (Fig. 4C and SFig. 7). The relative rates of these reactions also allowed us to attribute a predominant translocation state to the EC tested because nucleotide addition proceeded from post-translocation, pyrophosphorolysis from pre-translocation and hydrolysis from the backtracked state (scheme on Fig. 4C). Comparison of the rates of these reactions for the three complexes used in the present study suggested that EC14 is mainly stabilized in a post-translocated state (characterized by fast NTP addition), EC15 is mainly pre-translocated (fast pyrophospholysis), and EC16 is mainly backtracked/paused (faster hydrolysis), similar to the ECs formed by *Thermus aquaticus* RNAP (21), which doesn’t contain Si3, on this template. These results imply that Si3 does not influence the catalysis or translocation equilibrium of cyRNAP.

**Fig. 4.**
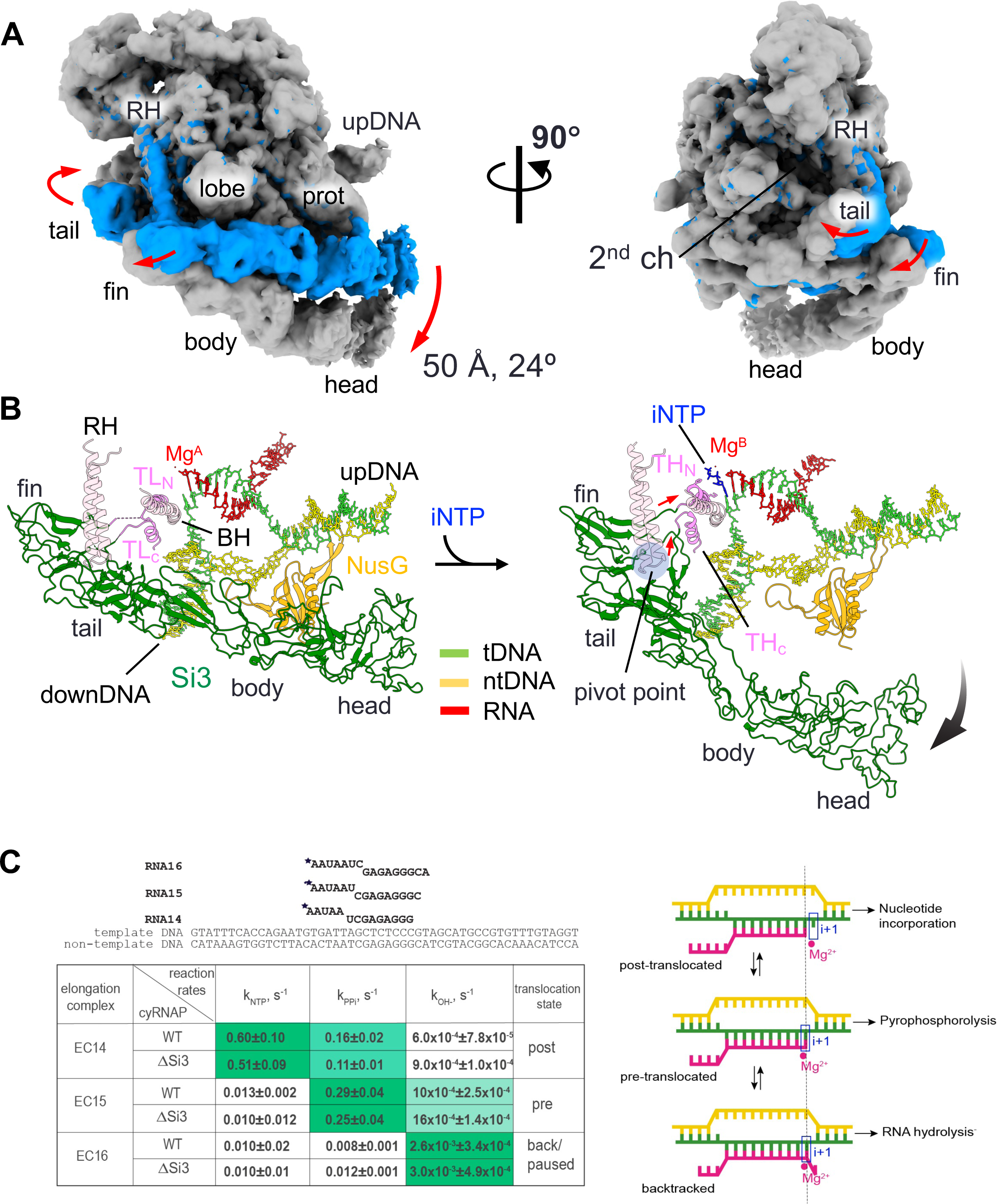
Si3 movement during the trigger helix folding. (A) Cryo-EM maps of the iNTP-bound (gray) and iNTP-free (light blue) states of the EC-NusG strains (RH, rim helix; prot, protrusion; upDNA, upstream DNA). Arrows indicate movement of Si3 by trigger helix folding. (B) Conformational change in Si3 during the transition from the trigger loop (TL) to the trigger helix (TH) by iNTP (blue stick model) binding. The red and black arrows indicate movements of the TL/TH-Si3 linker and Si3, respectively. A pivot point for converting the movement of the linker to the swing motion of Si3 is shown as a blue transparent circle. (C) Si3 does not influence catalysis by cyRNAP. Scheme and sequence of the assembled elongation complex used for experiments with WT and ΔSi3 RNAPs. The table represents the summary of reaction rate constants of single nucleotide addition (*k*_NTP_), pyrophosphorolysis (*k*_PPi_) and transcript hydrolysis (*k*_OH-_) in EC14, EC15 and EC16 by WT and ΔSi3 RNAPs. The values that follow the ± sign are the values of standard deviation derived from three independent experiments. The shade of green in the cells reflects the value of the constant, i.e., darkest shade corresponds to the highest rate. The right column shows the predominant translocation states of the elongation complexes, as deduced from the relative rates of reaction. Scheme of RNAP oscillation in translocation equilibrium and the architecture of the nucleic acid scaffold of the elongation complex in postLJtranslocation, preLJtranslocation and backtracked states, as adapted from (21). The template DNA, the non-template DNA and the RNA are green, yellow and pink, respectively. Catalytic Mg^2+^ ions and the i+1 site of the RNAP active center are shown by a red circle and a blue rectangle, respectively.

The cryo-EM structure of the cyRNAP-promoter DNA complex containing σ^A^ (both from *Synechocystis* sp. PCC 6803, which is closely related to the *S. elongatus* PCC 7942 used in this study), promoter DNA and 4-mer RNA was determined by Shen et al. (15); the results showed that Si3-head contacts σ^A^ domain 2. This interaction clamps the single-stranded DNA around the -10 region, stabilizing the open complex and facilitating transcription initiation. Comparison of the structures of the cyRNAP promoter complex (15) with those of the EC (this study) revealed that Si3-body and -head move toward σ^A^ domain 2 for interaction but that the other cyRNAP structures, including Si3-tail and -fin and the main body of the RNAP, are nearly identical (Fig. 5A).

**Fig. 5.**
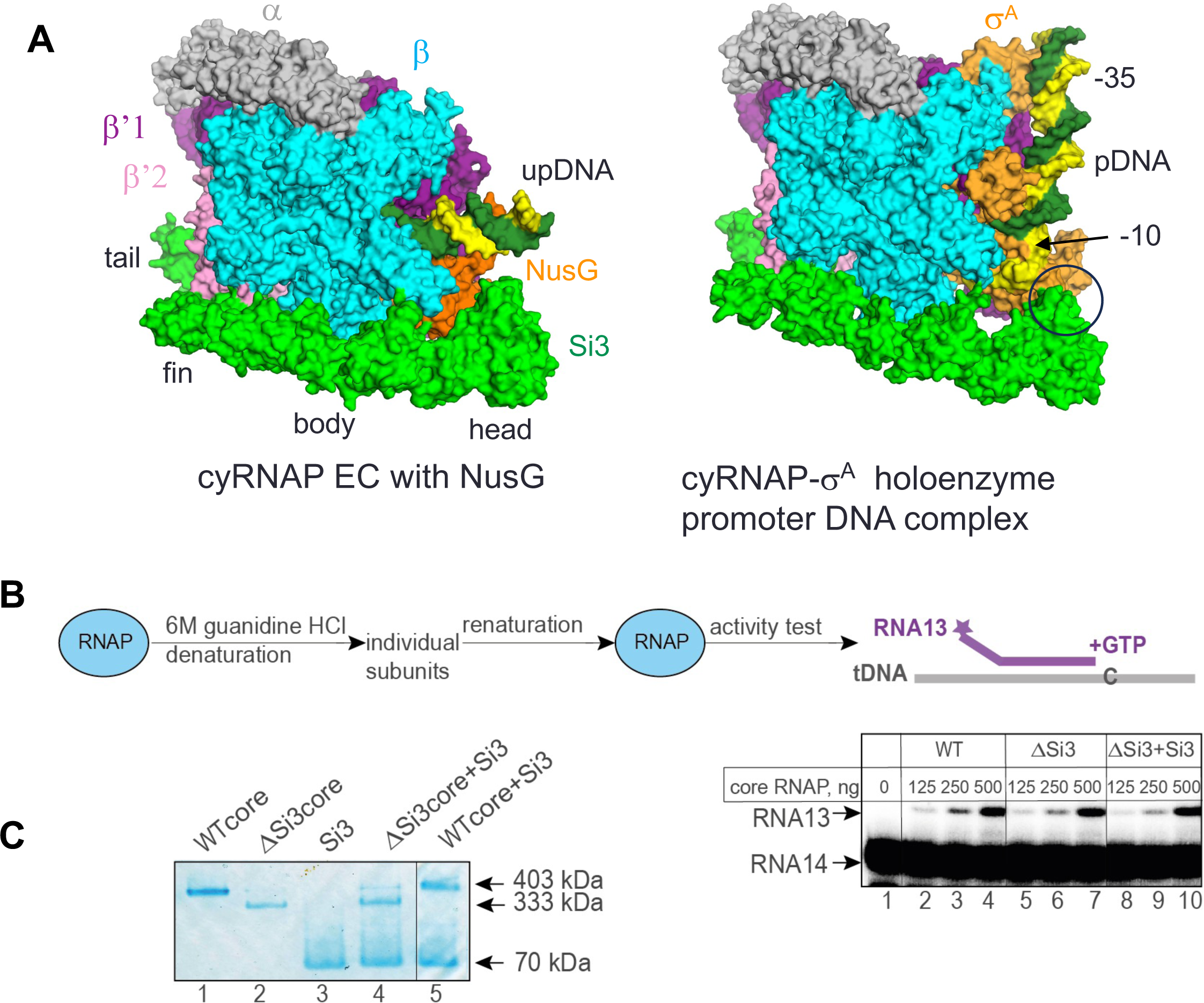
Si3 functions. (A) The cryo-EM structures of cyRNAP in the EC (left) and the promoter complex (right, PDB: 8GZG). The contact between Si3-head and σ^A^ in the promoter complex is indicated by a black circle. (B) Si3 is not required for cyRNAP assembly or maturation. WT and ΔSi3 cyRNAPs were denatured and subsequently renatured, after which their activity was tested on the construct mimicking the DNA template–RNA transcript duplex structure by their ability to incorporate the next nucleotide, G, dictated by the template. Twofold serial dilutions of the assembly mixture with the indicated initial amounts of core enzymes were tested. The vertical lines indicate the positions where the parts of the same gel were combined. (C) The recombinant Si3 protein can bind ΔSi3 cyRNAP but not WT cyRNAP. The complex formation between the indicated proteins was analyzed by blue native polyacrylamide gel electrophoresis. The vertical line indicates the position where two parts of the same gel were combined.

Si3 wraps around the main body of cyRNAP, which may facilitate RNAP folding, subunit assembly and/or maturation to form an active and mature form of RNAP as DNA and a σ factor that enhances reconstitution of *E. coli* RNAP (22). To test the function of Si3 during cyRNAP assembly and maturation, we performed a refolding experiment with WT, ΔSi3 cyRNAP and ΔSi3 cyRNAP in combination with the separately expressed and purified Si3 protein (ΔSi3+Si3) (Fig. 5B). The proteins were denatured with 6 M guanidine-HCl and renatured by gradual removal of guanidine-HCl via dialysis against renaturation buffer. The activities of the reconstituted ΔSi3 cyRNAP in the absence and presence of the Si3 protein, as judged by their ability to extend 13 nt long RNA in the assembled duplex with template DNA oligonucleotide, were nearly the same as those of the WT cyRNAP, indicating that Si3 does not play a role in cyRNAP assembly and maturation. This conclusion is supported by the similar yields of recombinant WT and ΔSi3 cyRNAPs routinely isolated from *E. coli.* Remarkably, however, the separate Si3 protein binds ΔSi3 cyRNAP but not the WT cyRNAP when it is added externally to cyRNAP (Fig. 5C). When complex formation between Si3 and ΔSi3 cyRNAP was assessed by a blue native polyacrylamide gel electrophoresis, a band with a lower mobility similar to that of the WT cyRNAP was observed (Fig. 5C, Lane 4). Interaction between WT cyRNAP and Si3 was not detected, i.e., no complex with lower mobility than that of WT cyRNAP was detected (Lane 5).

## Discussion

In this study, we determined the structures of cyRNAP Si3 by X-ray crystallography (Fig. 1) and of cyRNAP EC-NusG with and without iNTP by cryo-EM (Figs. 2 and 4). We investigated the function of Si3 by comparing the catalytic activities of WT and ΔSi3 cyRNAPs. The results of structural and biochemical investigations of cyRNAP showed that Si3 is accommodated within the cavities of cyRNAP without compromising its basic activities, that it shields the site of secondary channel binding proteins, and that it moves within cyRNAP upon binding of iNTP in the active site. Remarkably, a minor structural transition between the trigger loop and trigger helix causes a major swinging motion of Si3 (Fig. 4 and SMovie 2). The presence of Si3 in the middle of the trigger loop/helix did not affect cyRNAP catalysis under our experimental conditions (Fig. 4C). Because of the large conformational change that occurs during the transcription reaction, changes in cyRNAP activity could be observed when the motion of Si3 is hindered, such as by binding of external factors. Further proteomics for searching factors binding Si3, structural, single-molecule and biochemical studies are required to elucidate its role in regulating transcription by cyRNAP, such as by sensing environmental signals (e.g., trafficking of RNAP or transcription-translation coupling) to optimizing cyRNAP activity. Alternatively, the oscillating motion of Si3 might function as a regulatory signal for cellular processes. Photosynthetic cyanobacteria synchronize their gene expression patterns with diurnal light cycles (23). Conceivably, the lack of Si3 movement might trigger initiation of cyRNAP hibernation through binding to cellular factors or its oligomerization during the night. Additionally, Si3 movement might help RNAP propel through the densely packed cytoplasm of cyanobacteria during transcription.

The primary proofreading mechanism employed by RNAP involves backtracking followed by hydrolysis of misincorporated nucleotides at the 3’-end of nascent RNA. This process is significantly enhanced by elongation factors that bind to RNAP secondary channel, such as Gre in bacteria, TFS in archaea, and TFIIS in eukaryotes (24). However, unlike the absolute majority of living organisms, cyanobacteria lack Gre factor. The intracellular concentration of Mn^2+^ is two orders of magnitude greater in cyanobacteria than in other bacteria to support photosynthesis (16). It is possible that Mn^2+^ replaces the catalytic Mg^2+^ of RNAP and thus promotes misincorporation of NTPs (25, 26). Potentially as a compensating mechanism, cyRNAP has been shown to possess proficient intrinsic proofreading activity (7, 27). However, this intrinsic activity is still approximately 10 times lower than the Gre-stimulated activity of *E*.

*coli* RNAP. Gre-like factors either emerged after the split of cyanobacteria from their last common ancestor with other bacteria or were subsequently lost. The distinctive characteristics of cyRNAP—the absence of Gre/DksA factors and the split of the largest subunit may be intrinsically linked to Si3 acquisition. The Si3-tail/fin position around the rim helix of RNAP prevents association of secondary channel binding proteins, such as GreA and DksA, with cyRNAP (Fig. 3). As secondary channel binding proteins play critical roles in transcription fidelity and regulation in bacteria, the Si3-GreA/DksA trade-off in cyanobacteria might be advantageous but remains to be fully understood. With Si3 acquisition, β’ increased to 210 kDa in size, and separation of the original *rpoC* gene into two genes was perhaps beneficial to facilitate expression of such a large protein. The observed change in the position and mobility of Si3 in cyRNAP ECs compared to those in the promoter complex (Fig. 5A) raises questions about the role of Si3 in promoter escape. Si3 may complicate promoter escape by binding to the σ factor; conversely, its large-range movement upon RNA synthesis may contribute to weakening σ association with core and/or promote σ release at transition to elongation stage.

The structure corresponding to Si3 of cyRNAP has not been found in other bacterial RNAPs. However, the structure and arrangement of the Rpb9 subunit in eukaryotic RNAPII show remarkable similarity to those of the Si3 subunit of cyRNAP (Fig. 3C). Rpb9 is positioned within a cavity between the rim helix and the lobe domain of RNAPII, akin to the Si3-fin of cyRNAP (highlighted in red in cyRNAP and RNAPII). Rpb9 is a unique subunit found only in RNAPII and plays a critical role in enhancing the accuracy of transcription (28). Although both Rpb9 and Si3-tail are located away from the active site of RNAP, their presence may enhance transcription fidelity, which coordinates RNAP confirmation changes such as RNAP swiveling and/or movement of the rim helix during the nucleotide addition cycle (20). The presence of these unique structural features in different types of RNAPs suggests a common mechanism for enhancing transcriptional accuracy and specificity across different organisms. Further investigation of Si3 function at different stages of transcription and under several growth conditions in cyanobacteria will be required to determine the full array of its biological functions.

## Experimental Procedures

### Protein preparation

The DNA fragment encoding *Thermosynechococcus elongatus* BP-1 Si3 in the β’2 subunit (TelSi3, RpoC2 residues 345-983, 69 kDa) was cloned between the NdeI and BamHI sites of the pET15b expression vector to introduce an N-terminal His_6_-tag, and the protein was overexpressed in *E. coli* BL21(DE3)/pLysS cells. Transformants were subsequently grown in LB media supplemented with ampicillin (100 μg/ml) and chloramphenicol (25 μg/ml) at 37 °C until the OD_600_ reached ∼0.5, after which protein expression was induced by adding 0.5 mM IPTG for 10 h at 4 °C. The harvested cells were lysed by sonication, and proteins in the soluble fraction were purified by Ni-affinity column chromatography (HisTrap 5 ml column, GE Healthcare). The His_6_-tag was removed by thrombin digestion (1 μg of thrombin per mg of TelSi3 protein) for 20 h at 4 °C, and the protein was further purified by Q Sepharose column chromatography (GE Healthcare) and gel-filtration column chromatography (HiLoad Superdex75 16/60, GE Healthcare). The purified protein was concentrated to 15 mg/ml and exchanged into buffer containing 10 mM Tris-HCl (pH 8.0), 50 mM NaCl and 0.1 mM EDTA.

### Limited trypsinolysis

Limited trypsinolysis was used to remove flexible regions from TelSi3, and N-terminal amino acid sequencing was used to identify protein fragments suitable for crystallization. The trypsin digests were carried out in 10 mM Tris–HCl (pH 8), 100 mM NaCl, 5% (v/v) glycerol, 0.1 mM EDTA and 1 mM DTT. TelSi3 (10 mg/ml) was digested in a 10 µl volume with different amounts of trypsin (5 nM to 5 µM) for 10 min at 25 °C. The reactions were terminated by addition of PMSF. The trypsinized fragments were separated by SDSLJPAGE and blotted onto PVDF membranes, and the N-terminal sequences were determined by Edman based protein sequencing. The TelSi3 fragment containing residues 435-938 (TelSi3ΔN, 60 kDa) was PCR subcloned and inserted into the pET15b expression vector between the NdeI and BamHI sites. The protein was overexpressed and purified as described above for full-length TelSi3.

### Crystallization

Initial crystals of TelSi3ΔN were obtained by the hanging-drop vapor diffusion method by mixing equal volumes of the protein solution (20 mg/ml) and crystallization solution (0.1 M sodium citrate [pH 3.5], 0.2 M MgCl_2_ and 10% PEG6000) and incubating at 4 °C over the same crystallization solution. The large crystals (0.5 × 0.2 × 0.2 mm) used for X-ray data collection were prepared by microseeding by mixing 2 µl of protein solution, 2 µl of crystallization solution (0.1 M sodium citrate [pH 5], 0.4∼0.6 M MgCl_2_, 4∼6% PEG3350 and 50 µg/ml heparin) and 0.2 µl of seed solution. The crystals were then dehydrated by transfer to crystallization solution (without heparin) with increasing concentrations of PEG3350 (in 5% steps) to a final concentration of 20% and incubated for 5-10 h. For all procedures, crystal preparation, growth and dehydration were performed at 4 °C. The crystals were transferred to a crystallization solution with 25% (v/v) propylene glycol as a cryoprotective solution and flash frozen in liquid nitrogen. Selenomethionine-substituted proteins were prepared for SAD analysis by suppressing methionine biosynthesis.

### X-ray data collection and crystal structure determination

In addition to the four original methionine residues found in TelSi3ΔN (including an N-terminal methionine residue resulting from cloning into the pET15b vector), three methionine residues were introduced by replacing the leucine residues at 508, 738 and 922 by site-directed mutagenesis to obtain the experimental phase via single-anomalous dispersion (SAD) experiments using SeMet-labeled proteins. The TelSi3ΔN protein with seven selenomethionine residues (TelSi3ΔN^Met7^) was generated by suppressing methionine biosynthesis during overexpression of the TelSi3ΔN^Met7^ protein. The protein was purified as described above.

Diffraction data were corrected at National Synchrotron Light Source (NSLS) beamline X25. There are six TelSi3ΔN^Met7^ molecules in an asymmetric unit of the crystal belonging to the P3(2)21 space group. The crystallographic datasets were processed using HKL2000 (29). With the anomalous signal from SeMet, the experimental phase (figure of merit: 0.273) was calculated using automated structure solution (AutoSol) in PHENIX (30). Density modification yielded a map suitable for manual model building by Coot (31) followed by structure refinement using PHENIX. The final coordinates and structure factors have been deposited in Protein Data Bank (PDB) under the accession codes listed in Supplementary Table 1.

### Expression and isolation of the *Synechococcus elongatus* RNAP

The core enzyme of cyRNAP was overexpressed in *E. coli* T7Express cells (New England Biolabs) cells transformed with a pET28a expression vector containing the α, β, β’1, β’2 and ω encoding genes (β and β’2 contain a Strep-tag and His-tag, respectively) (32). The cells were grown in LB media supplemented with kanamycin (50 μg/ml) at 37 °C until the OD_600_ was ∼0.6. Afterward, the cells were induced with IPTG (1 mM) and grown overnight at 22 °C. The biomass was harvested and suspended in lysis buffer (50 mM Tris-HCl (pH 8.0), 250 mM NaCl, 10% glycerol, 20 mM imidazole, and 1 mM β-mercaptoethanol and protease inhibitors from Roche according to the manufacturer’s instructions). The cells were sonicated, lysate centrifuged at 18 k x g, after which the supernatant was collected. The protein was purified at 4 °C sequentially through a HisTrap (5 mL) column and a Strep-Tactin XT (1 mL) column (both from Cytiva). The latter column was washed with 3 column volumes (CVs) of Buffer W (100 mM Tris-HCl pH 8.0, 150 mM NaCl, 1 mM EDTA). The bound protein was eluted by applying 1 CV of Buffer E (100 mM Tris-HCl pH 8.0, 150 mM NaCl, 1 mM EDTA, 2.5 mM desthiobiotin). The purified cyRNAP (20 μM) was assessed using SDSLJPAGE, dialyzed against Storage Buffer (40 mM Tris–HCl pH 8.0, 200 mM KCl, 1 mM EDTA, 1 mM DTT, and 5% glycerol), and stored at -80 °C.

### Cloning, expression and isolation of the *Synechococcus elongatus* NusG and Si3 proteins

The NusG was overexpressed in *E. coli* T7Express cells (New England Biolabs) cells transformed with pET28a expression vector where the gene for the C-terminal His_6_-tagged *Synechococcus elongatus* NusG was cloned. Cells were grown in LB medium supplemented with kanamycin (50 μg/ml) at 37°C until OD_600_ ∼0.5, then induced with IPTG (1 mM) and grown overnight at 22°C. Culture pellets were sonicated in 50 ml Lysis Buffer (10 mM Tris-HCl pH 7.9, 300 mM KCl, protease inhibitors from Roche according to manufacturer), spun at 18K rpm, and filtered through 0.22 μM syringe filter. Filtered supernatant was subjected to Ni-NTA affinity chromatography in 10 mM Tris-HCl pH 7.9, 600 mM KCl, 5% glycerol with 50 mM imidazole washes and 100 mM imidazole elution. The eluted protein (in 600 mM KCl) was diluted (∼100 mM KCl) and applied to a pre-equilibrated with (10 mM Tris-HCl pH 8.0, 100 mM KCl, 5% glycerol) 5 ml Resource Q column, Cytiva. The column was washed with 5 CV of equilibration buffer, and the protein was eluted by applying a linear salt gradient (100-1 M KCl) over 10 CV. The purified NusG (90 μM) was checked using SDS-PAGE, stored in Storage Buffer (40 mM Tris-HCl pH 8.0, 200 mM KCl, 1 mM EDTA, 1 mM DTT, 5% glycerol) at -80 °C. Cyanobacteria-specific loop of NusG (residues 110-122) was deleted by site-directed mutageneses, and the mutant NusG was isolated as the WT protein. The open reading frame encoding separate full-size Si3 domain was cloned into pET28 vector, overexpressed *E. coli* T7Express cells (New England Biolabs) as the N-terminal His_6_-tagged protein, and isolated via Ni-NTA affinity chromatography on HisTrap column, Cytiva, similarly to NusG protein. After affinity chromatography the protein was dyalised against the storage buffer (20 Tris-HCl pH 8.0, 200 mM KCl, 1 mM EDTA, 1 mM DTT, 50% glycerol).

### Sample preparation for cryo-EM

The cyRNAP EC with NusG was reconstituted in vitro by mixing 5 μM cyRNAP with equimolar amounts of template DNA and RNA (Fig. 2A) in storage buffer at 37 °C for 10 minutes, followed by mixing with 7 μM nontemplate DNA and incubating further for 10 minutes. The resulting EC was mixed with 7 μM NusG and incubated for 10 min at 37 °C. CHAPSO (8 mM) was added to the sample just before vitrification. The iNTP-bound EC was prepared by adding 1 mM 3’-deoxyATP or CTP to the EC with NusG and incubating for 5 min at 37 °C. Another difference between the EC- and iNTP-bound ECs is the nontemplate DNA used in the scaffold, the latter of which contains complementary transcription bubbles.

### Grid preparation for cryo-EM

C-flat Cu grids (CF-1.2/1.3 400 mesh, Protochips, Morrisville, NC) were glow-discharged for 40 seconds using the PELCO easiGlow^TM^ system prior to application of 3.5 μl of the sample (2.5 – 3.0 mg/ml protein concentration) and plunge-freezing in liquid ethane using a Vitrobot Mark IV (FEI, Hillsboro, OR) with 100% chamber humidity at 5 °C.

### Cryo-EM data acquisition and processing

Data were collected using a Titan Krios (Thermo Fisher) microscope equipped with a Falcon IV direct electron detector (Gatan) at Penn State Cryo-EM Facility. Sample grids were imaged at 300 kV, with an intended defocus range of -2.5 to -0.75 μm and a magnification of 75,000X in electron counting mode (0.87 Å per pixel). Movies were collected with a total dose of 45 electrons/Å^2^. Downstream processing was performed with CryoSPARC (33). The movies were corrected and aligned using patch motion correction followed by patch CTF correction. Particles were picked using a template-based autopicker and multiple rounds of 2D classification to discard bad particles. The 2D classes with EC-NusG particles were selected and used for training the Topaz model (34). The Topaz-extracted particles were subjected to multiple rounds of heterogeneous refinement to remove junk particles. Finally, a nonuniform refinement operation was run on the final set of particles to yield the reconstruction (SFigs. 4 and 6).

### Structure refinement and model building

A model of the cyRNAP core enzyme was constructed by homology modeling using core RNAP from the cryo-EM structure of the *Syn*6803 RNAP-σ^A^ promoter DNA open complex as a reference model. A model of cyNusG was constructed with the AlphaFold2 gene (35). DNA and RNA models were constructed using the *E. coli* RNAP elongation complex (PDB: 7MKO) as a guide. The cyRNAP gene was manually fitted into the cryo-EM density map using Chimera (36), followed by rigid body and real-space refinement using Coot (31) and Phenix (37).

### *In vitro* transcription in the assembled elongation complexes

ECs were assembled and immobilized as described (38). Sequences of the oligonucleotides used for the assembly of ECs are shown on Fig.4C. For assembly of ECs used for experiments on Fig. 4C, 13 nt long RNA was radiolabelled at the 5’-end with [γ-^32^P] ATP and T4 Polynucleotide kinase (New England Biolabs) prior to complexes assembly. Stalled elongation complexes EC14, EC15 and EC16 were obtained by extension of the initial RNA13 in EC13 with 10 μM NTP sets according to the sequence for 5 min and then were washed with TB to remove Mg^2+^ and NTPs. Reactions were initiated by addition of 10 mM MgCl_2_ with or without either 1 μM NTPs or 250 μM PPi. Single nucleotide addition and pyrophosphorolysis experiments were performed at 30°C in transcription buffer (TB) containing 20 mM Tris–HCl pH 6.8, 40 mM KCl, 10 mM MgCl_2_, transcript hydrolysis was done in the same buffer except at pH 7.9. After incubation for intervals of time specified on Figures, reactions were stopped with formamide-containing buffer. Products were resolved by denaturing 23% polyacrylamide gel electrophoresis (PAGE) (8 M Urea), revealed by PhosphorImaging (Cytiva) and visualized using ImageQuant (Cytiva) software. Kinetics data were fitted to a single exponential equation y=y_0_+a^-bx^ using SigmaPlot software by non-linear regression to determine rate constants of the reactions.

### Denaturation and renaturation of cyRNAP

Denaturation of cyRNAP was performed by incubating the purified protein for 20 min in denaturing buffer containing 20 mM Tris-HCl (pH 7.9), 6 M guanidine-HCl, 5% glycerol, 1 mM EDTA, and 10 mM DTT at 30 °C in a 100 µl volume and with a cyRNAP concentration of 0.5 mg/ml. Recombinant Si3 was included in 2.5 molar excess. The proteins were renatured via overnight dialysis at 7 °C against renaturing buffer containing 20 mM Tris-HCl (pH 7.9), 200 mM KCl, 10% glycerol, 2 mM MgCl_2_, 10 µM ZnCl_2_, 1 mM EDTA, and 1 mM DTT. Aliquots of the renaturation mixture and their serial dilutions were used for nucleotide addition experiments on assembled constructs containing template DNA and RNA oligonucleotides. A 13 nt RNA oligonucleotide was radiolabeled at the 5’ end with [γ-^32^P] ATP and T4 polynucleotide kinase (New England Biolabs) prior to EC assembly. The indicated on the Fig. 5C amount of assembly # mixture was incubated with the RNA-DNA duplex for 5 min at room temperature, then 10 µM GTP was added for 10 minutes at 30°C. Reactions were stopped and products analyzed as before.

### Complex formation between the Si3 protein and core cyRNAP

For the binding experiment 150 nM core enzymes and 1.5 µM Si3 proteins were incubated for 10 minutes at 4°C in 20 mM Tris-HCl pH 7.9, 40 mM KCl, mixed with loading dye (final concentration is 50mM BisTris pH 7.2, 50mM NaCl, 10% glycerol, 0.001% Ponceau S) and resolved on the NativePAGE 3-12% Bis-Tris gel, Invitrogen using running buffers prepared according to the manufacturer, for 90 minutes at 150V. Gel was fixed with 50% methanol,10% acetic acid solution, and additionally de-stained by boiling in 8% acetic acid.

### Salt stability of elongation complexes

Elongation complex was assembled using oligos shown on SFig. 5B. 14 nt RNA in ECs on was radiolabelled at the 3’ end by incorporation of [α-^32^P] GTP into original 13 nt long RNA. To examine the stability of ECs, ECs bound to the streptavidin sepharose beads, Cytiva via strep tag on β subunit of cyRNAP, were incubated in TB containing 300 mM KCl at 30°C for times specified on SFig. 5B. WT or mutant NusG^Δ110-122^ were added where specified at 1 μM final concentration. Supernatant and total fractions were collected for analysis. Reactions were stopped and products analyzed as before.

### Data, Materials, and Software Availability

The X-ray crystallographic density map and the refined model have been deposited in Protein Data Bank (www.rcsb.org) under accession number 8EMB. The cryo-EM density map and the refined model have been deposited in Electron Microscopy Data Bank (www.ebi.ac.uk/emdb/) under accession numbers EMD-40874 (iNTP-free EC-NusG) and EMD-42502 (iNTP-bound EC-NusG) and in Protein Data Bank (www.rcsb.org) under accession numbers 8SYI (iNTP-free EC-NusG) and 8URW (iNTP-bound EC-NusG). All study data are included in the article and/or SI Appendix.

## Supporting information

SMovie 1

SMovie 2

## ACKNOWLEDGMENTS

We thank Jean-Paul Armache at Penn State for the technical support. We thank the National Synchrotron Light Source (NSLS) Brookhaven National Laboratory for X-ray data collection. We would like to acknowledge the Penn State Huck Life Science Institutes Cryo-EM Core Facility for use of the Talos Arctica G2 TEM and the Vitrobot Mark IV and Sung Hyun Cho for data collection. We thank Yu Zhang at the Shanghai Institute of Plant Physiology and Ecology for kindly sharing the coordinates of *Synechocystis* sp. PCC 6803 RNAP. This work was supported by a National Institutes of Health grant (R35 GM131860 to K. S. M.) and a Biotechnology and Biological Sciences Research Council grant BB/W017385/1 to Y.Y.

