## Supplementary material for "Structure and function of the Si3 insertion integrated into the trigger loop/helix of cyanobacterial RNA polymerase": SI Figures

### SFig. 1

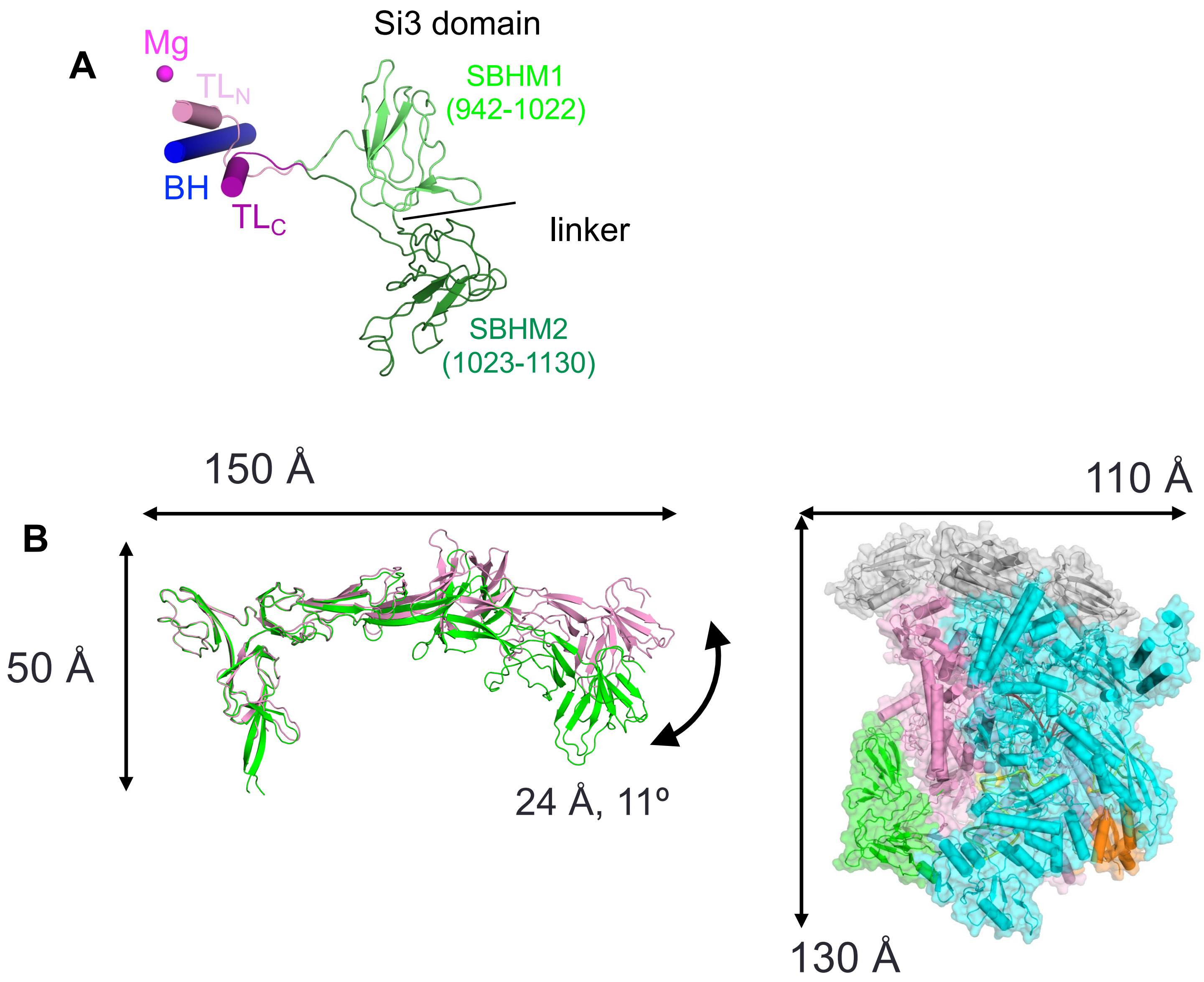


**SFig. 1**. (A) Structure of the Si3 domain (light and dark green) of *E. coli* RNAP, its connection to the trigger loop (TL). $Mg^{2+}$ bound at the active site and bridge helix (BH) are depicted as a sphere and cartoon models. (C) Comparing the sizes of Si3 domain (without Tail sub-domain, SBHM1) of cyanobacterial RNAP and the *E. coli* RNAP (Si3 of the *E. coli* RNAP is highlighted in green).

SFig. 2

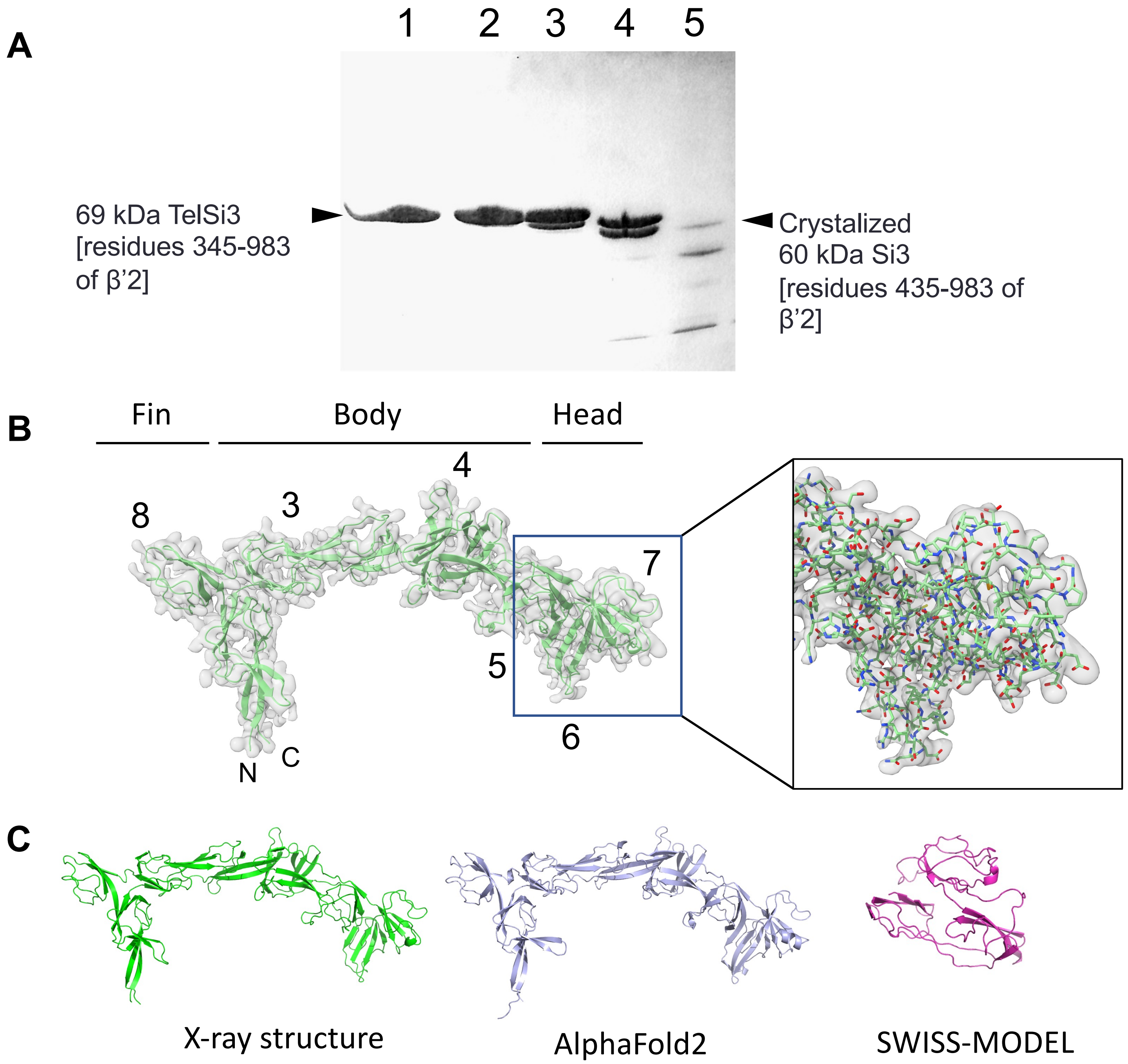


**SFig. 2**. (A) Limited trypsinolysis of the TelSi3 protein analyzed by SDS–PAGE. Reactions contained 10 mg/ml of TelSi3 and 0, 5, 50, 500 and 5000 nM of trypsin (lanes 1 to 5, respectively). (B) 2*Fo-Fc* electron density map of the TelSi3dΔN (gray transparent, σ cutoff=1.3) overlaid on a protomer I in an asymmetric unit (green cartoon model). The SBHM are labeled (2 to 8) and sub-domains are indicated. Magnified view of a boxed area is shown with a stick model of the TelSi3dΔN (Head sub-domain). (C) Comparing the X-ray structure of TelSi3dΔN (left) with the structures predicted by AlphaFold2 (middle) and SWISS-MODEL (right).

SFig. 3

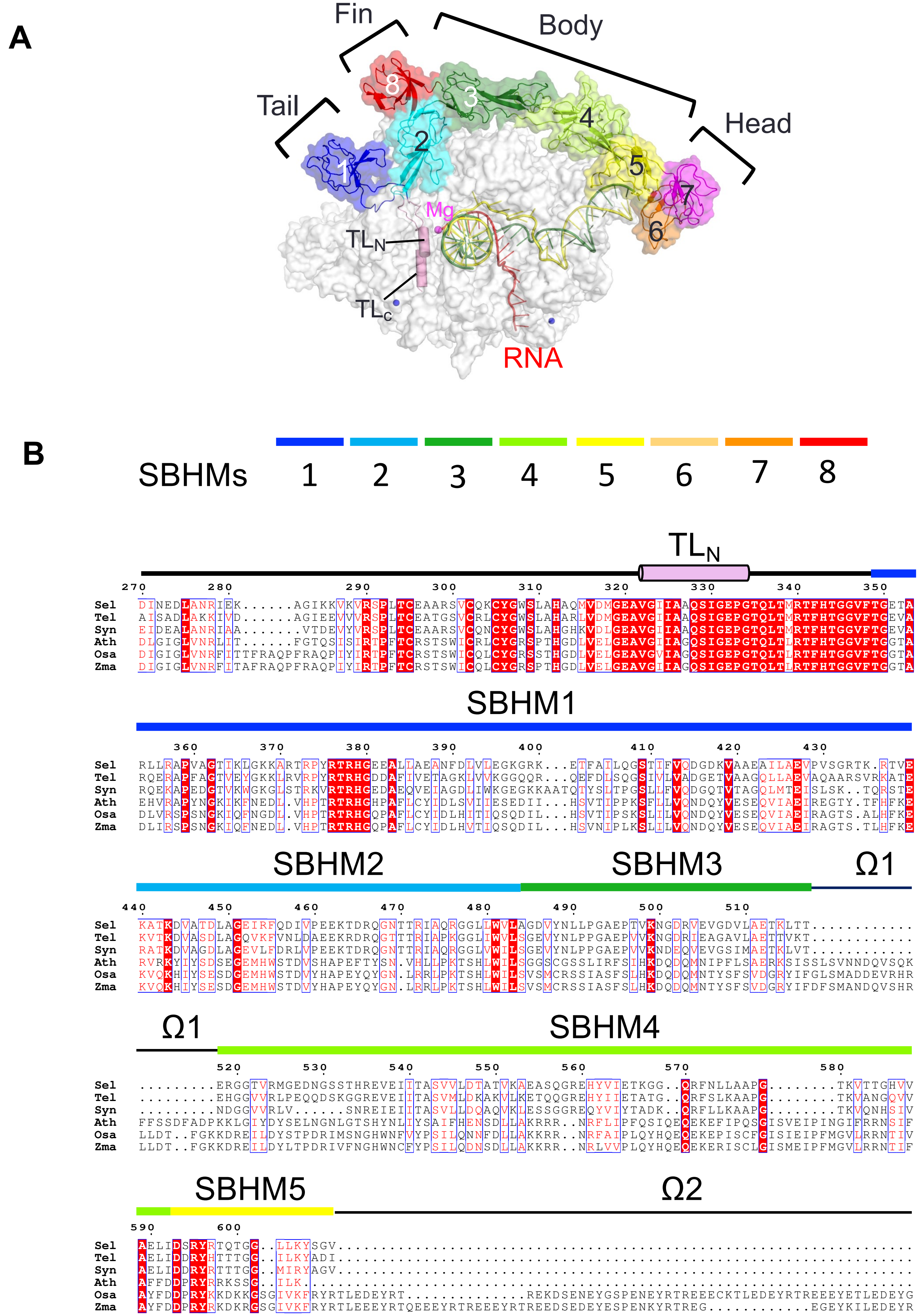

A
Fin
Body
Tail
Head
1
2
3
4
5
6
7
8
Mg
TLN
TLC
RNA
B
SBHMs 1 2 3 4 5 6 7 8
TLN
270 280 290 300 310 320 330 340 350
Sel DINEDLANRIEK......AGIKKVKVRSPLTCEAARSVCQKCYGWSLAHAQMVDMGEAVGIIAAQSIGEPGTQLTMRTFHTGGVFTGETA
Tel AISADLAKKIVD......AGIEEVVVRSPLTCEATGSVCRLCYGWSLAHARLVDMGEAVGIIAAQSIGEPGTQLTMRTFHTGGVFTGEVA
Syn EIDEALANRIAA.......VTDEVYVRSPLTCEAARSVCQNCYGWSLAHGHKVDLGEAVGIIAAQSIGEPGTQLTMRTFHTGGVFTGEVA
Ath DLGIGLVNRLIT......FGTQSISIRTPFTCRSTSWICRLCYGRSPTHGDLVELGEAVGIIAGQSIGEPGTQLTLRTFHTGGVFTGGTA
Osa DIGIGLVNRFITTFRAQPFRAQPIYIRTPFTCRSTSWICQLCYGRSSTHGDLVELGEAVGVIAGQSIGEPGTQLTLRTFHTGGVFTGGTA
Zma DIGIGLVNRFITAFRAQPFRAQPIYIRTPFTCRSTSWICQLCYGRSPTHGDLVELGEAVGIIAGQSIGEPGTQLTLRTFHTGGVFTGGTA
SBHM1
360 370 380 390 400 410 420 430
Sel RLLRAPVAGTIKLGKKARTRPYRTRHGEEALLAEANFDLVLEGKGRK...ETFAILQGSTIFVQDGDKVAAEAILAEVPVSGRTK.RTVE
Tel RQERAPFAGTVEYGKKLRVRPYRTRHGDDAFIVETAGKLVVKGGQQR...QEFDLSQGSIVLVADGETVAAGQLLAEVAQAARSVRKATE
Syn RQEKAPEDGTVKWGKGLSTRKVRTRHGEDAEQVEIAGDLIWKGEGKKAATQTYSLTPGSLLFVQDGQTVTAGQLMTEISLSK..TQRSTE
Ath EHVRAPYNGKIKFNEDL.VHPTRTRHGHPAFLCYIDLSVIIESEDII...HSVTIPPKSFLLVQNDQYVESEQVIAEIREGTY.TFHFKE
Osa DLVRSPSNGKIQFNGDL.VHPTRTRHGQPAFLCYIDLHITIQSQDIL...HSVTIPSKSLILVQNDQYVESEQVIAEIRAGTS.ALHFKE
Zma DLIRSPSNGKIQFNEDL.VHPTRTRHGQPAFLCYIDLHVTIQSQDIL...HSVNIPLKSLILVQNDQYVESEQVIAEIRAGTS.TLHFKE
SBHM2
SBHM3
Ω1
440 450 460 470 480 490 500 510
Sel KATKDVATDLAGEIRFQDIVPEEKTDRQGNTTRIAQRGGLLWVLAGDVYNLLPGAEPTVKNGDRVEVGDVLAETKLTT............
Tel KVTKDVASDLAGQVKFVNLDAEEKRDRQGTTTRIAPKGGLIWVLSGEVYNLPPGAEPVVKNGDRIEAGAVLAETTVKT............
Syn RATKDVAGDLAGEVLFDRLVPEEKTDRQGNTTRIAQRGGLLWILSGEVYNLPPGAEPVVKNDEQVEVGSIMAETKLVT............
Ath RVRKYIYSDSEGEMHWSTDVSHAPEFTYSN.VHLLPKTSHLWILSGGSCGSSLIRFSIHKDQDQMNIPFLSAERKSISSLSVNNDQVSQK
Osa KVQKHIYSESDGEMHWSTDVYHAPEYQYGN.LRRLPKTSHLWILSVSMCRSSIASFSLHKDQDQMNTYSFSVDGRYIFGLSMADDEVRHR
Zma KVQKHIYSESDGEMHWSTDVYHAPEYQYGN.LRRLPKTSHLWILSVSMCRSSIASFSLHKDQDQMNTYSFSVDGRYIFDFSMANDQVSHR
Ω1
SBHM4
520 530 540 550 560 570 580
Sel .........ERGGTVRMGEDNGSSTHREVEIITASVVLDTATVKAEASQGREHYVIETKGG..QRFNLLAAPG........TKVTTGHVV
Tel .........EHGGVVRLPEQQDSKGGREVEIITASVMLDKAKVLKETQQGREHYIIETATG..QRFSLKAAPG........TKVANGQVV
Syn .........NDGGVVRLV......SNREIEIITASVLLDQAQVKLESSGGREQYVIYTADK..QRFLLKAAPG........TKVQNHSIV
Ath FFSSDFADPKKLGIYDYSELNGNLGTSHYNLIYSAIFHENSDLLAKRRR..NRFLIPFQSIQEQEKEFIPQSGISVEIPINGIFRRNSIF
Osa LLDT..FGKKDREILDYSTPDRIMSNGHWNFVYPSILQNNFDLLAKKRR..NRFAIPLQYHQEQEKEPISCFGISIEIPFMGVLRRNTIV
Zma LLDT..FGKKDREILDYLTPDRIVFNGHWNCFYPSILQDNSDLLAKKRR..NRLVVPLQYHQEQEKERISCLGISMEIPFMGVLRRNTIF
SBHM5
Ω2
590 600
Sel AELIDSRYRTQTGG..LLKYSGV.........................................................................
Tel AELIDDRYHTTTGG..ILKYADI.........................................................................
Syn AELIDDRYRTTTGG..MIRYAGV.........................................................................
Ath AFFDDPRYRKSSG..ILK..............................................................................
Osa AYFDDPRYKKDKKGSGIVKFRYRTLEDEYRT..............RKEDSENEYGSPENEYRTREEECKTLEDEYRTREEEYETLEDEYG
Zma AYFDDPRYRKDKRGSGIVKFRYRTLEEEYRTQEEEYRTREEEYRTREEDSEDEYESPENKYRTREG...........EGEYKILEDEYR

### SFig. 3

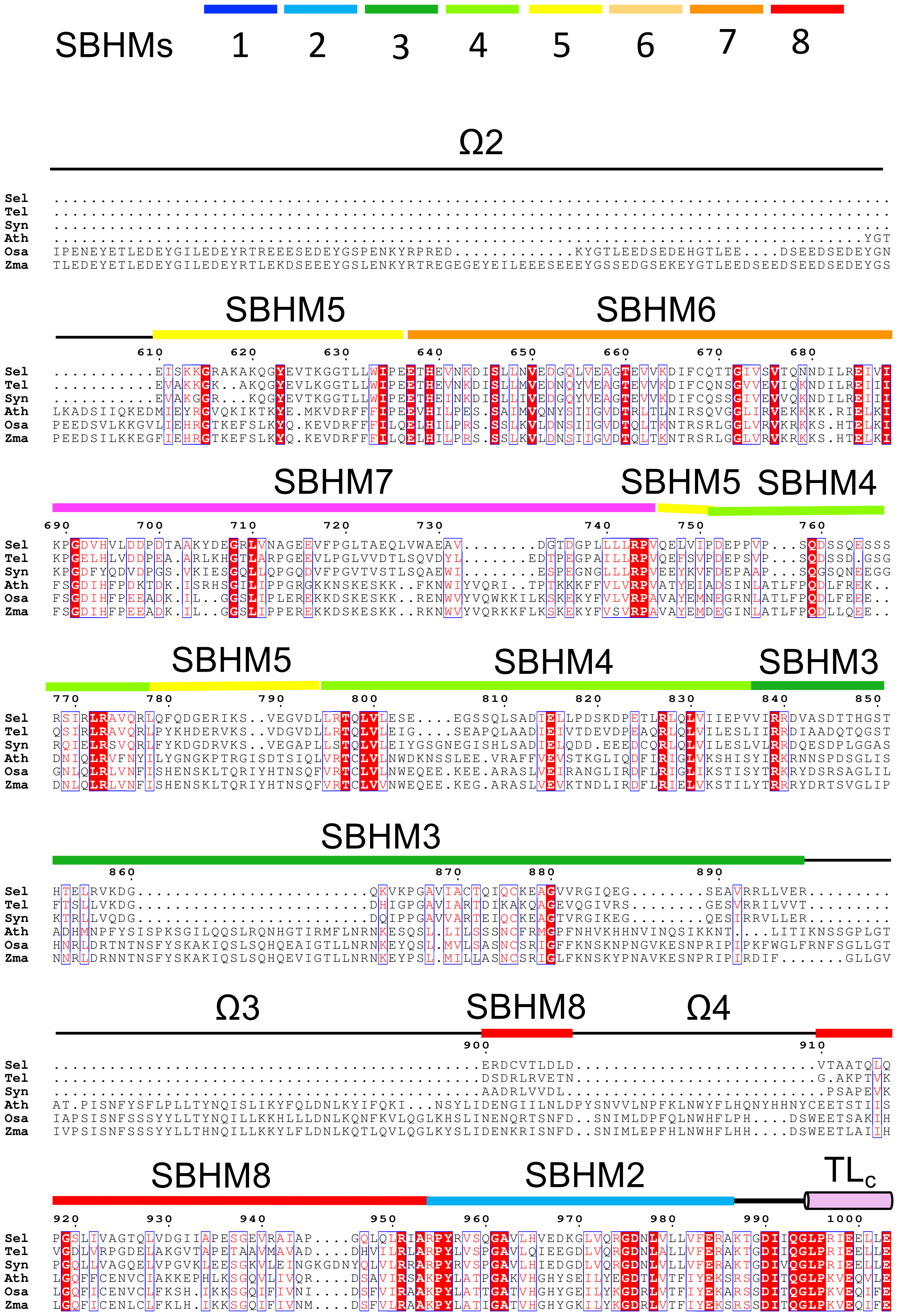

SBHMs 1 2 3 4 5 6 7 8
Ω2
Sel ............................................................................................
Tel ............................................................................................
Syn ............................................................................................
Ath .........................................................................................YGT
Osa IPENEYETLEDEYGILEDEYRTREEESEDEYGSPENKYRPRED...........KYGTLEEDSEDEHGTLEE....DSEEDSEDEYGN
Zma TLEDEYETLEDEYGILEDEYRTLEKDSEEEYGSLENKYRTREGEGEYEILEEESEEEYGSSEDGSEKEYGTLEEDSEEDSEEDSEDEYGS
SBHM5 SBHM6
610 620 630 640 650 660 670 680
Sel ..........EISKKGRAKAKQGYEVTKGGTLLWIPEETHEVNKDISLLNVEDGQLVEAGTEVVKDIFCQTTGIVSVTQNNDILREIVI
Tel ..........EVAKKGK..AKQGYEVLKGGTLLWIPEETHEVNKDISLLMVEDNQYVEAGTEVVKDIFCQNSGVVEVIQKNDILREIII
Syn ..........EVAKGGR...KQGYEVTKGGTLLWIPEETHEINKDISLLIVEDGQYVEAGTEVVKDIFCQSSGIVEVVQKNDILREIII
Ath LKADSIIQKEDMIEYRGVQKIKTKYE.MKVDRFFFIPEEVHILPES.SAIMVQNYSIIGVDTRLTLNIRSQVGGLIRVEKKKK.RIELKI
Osa PEEDSVLKKGVLIEHRGTKEFSLKYQ.KEVDRFFFILQELHILPRS.SSLKVLDNSIIGVDTQLTKNTRSRLGGLVRVKRKKS.HTELKI
Zma PEEDSILKKEGFIEHRGTKEFSLKYQ.KEVDRFFFILQELHILPRS.SSLKVLDNSIIGVDTQLTKNTRSRLGGLVRVKRKKS.HTELKI
SBHM7 SBHM5 SBHM4
690 700 710 720 730 740 750 760
Sel KPGDVHVLDDPDTAAKYDEGRLVNAGEEVFPGLTAEQLVWAEAV........DGTDGPLLLRPVQELVIPDEPPVP...SQDSSQESSS
Tel KPGELHLVDDPEA.ARLKHGTLARPGEEVLPGLVVDTLSQVDYL.......EDTPEGPAILLRPVQEFSVPDEPSVP...SQDSSD.GSG
Syn KPGDFYQDVDPGS.VKIESGQLLQPGQDVFPGVTVSTLSQAEWI........ESPEGNGLLLRPVEEYKVFDEPAAP...SQGSQNEEGG
Ath FSGDIHFPDKTDK.ISRHSGILIPPGRGKKNSKESKK..FKNWIYVQRI..TPTKKKFFVLVRPVATYEIADSINLATLFPQDLFREK..
Osa FSGDIHFPEEADK.IL..GGSLIPPEREKKDSKESKK..RENWVYVQWKKILKSKEKYFVLVRPAVAYEMNEGRNLATLFPQDLFEEE..
Zma FSGDIHFPEEADK.IL..GGSLIPPEREKKDSKESKK..RKNWVYVQRKKFLKSKEKYFVSVRPAVAYEMDEGINLATLFPQDLLQEE..
SBHM5 SBHM4 SBHM3
770 780 790 800 810 820 830 840 850
Sel RSIRLRAVQRLQFQDGERIKS..VEGVDLLRTQLVLESE....EGSSQLSADIELLPDSKDPETLRLQLVIIEPVVIRRDVASDTTHGST
Tel QSIRLRAVQRLPYKHDERVKS..VDGVDLLRTQLVLEIG....SEAPQLAADIEIVTDEVDPEAQRLQLVILESLIIRRDIAADQTQGST
Syn RQIELRSVQRLFYKDGDRVKS..VEGAPLLSTQLVLEIYGSGNEGISHLSADIELQDD.EEEDCQRLQLVILESLVLRRDQESDPLGGAS
Ath DNIQLRVFNYILYGNGKPTRGISDTSIQLVRTCLVLNWDKNSSLEE.VRAFFVEVSTKGLIQDFIRIGLVKSHISYIRKRNNSPDSGLIS
Osa GNLQLRLVNFISHENSKLTQRIYHTNSQFVRTCLVLNWEQEE.KEE.ARASLVEIRANGLIRDFLRIGLIKSTISYTRKRYDSRSAGLIL
Zma DNLQLRLVNFISHENSKLTQRIYHTNSQFVRTCLVVNWEQEE.KEG.ARASLVEVKTNDLIRDFLRIELVKSTILYTRRRYDRTSVGLIP
SBHM3
860 870 880 890
Sel HTELRVKDG........................QKVKPGAVIACTQIQCKEAGVVRGIQEG........SEAVRRLLVER.........
Tel FTSLLVKDG........................DHIGPGAVIARTDIKAKQAGEVQGIVRS........GESVRRILVVT.........
Syn KTRLLVQDG........................DQIPPGAVVARTEIQCKEAGTVRGIKEG........QESIRRVLLER.........
Ath ADHMNPFYSISPKSGILQQSLRQNHGTIRMFLNRNKESQSL.LILSSSNCFRMGPFNHVKHHNVINQSIKKNT....LITIKNSSGPLGT
Osa HNRLDRTNTNSFYSKAKIQSLSQHQEAIGTLLNRNKEYQSL.MVLSASNCSRIGFFKNSKNPNGVKESNPRIPIPKFWGLFRNFSGLLGT
Zma NNRLDRNNTNSFYSKAKIQSLSQHQEVIGTLLNRNKEYPSL.MILLASNCSRIGLFKNSKYPNAVKESNPRIPIRDIF.......GLLGV
Ω3 SBHM8 Ω4
900 910
Sel .............................................ERDCVTLDLD.........................VTAATQLQ
Tel .............................................DSDRLRVETN.........................G.AKPTVK
Syn .............................................AADRLVVDL...........................PSAPEVK
Ath AT.PISNFYSFLPLLTYNQISLIKYFQLDNLKYIFQKI...NSYLIDENGIILNLDPYSNVVLNPFKLNWYFLHQNYHHNYCEETSTIIS
Osa IAPSISNFSSSYYLLTYNQILLKKHLLFLDNLKQNFKVLQGLKHSLINENQRTSNFD..SNIMLDPFQLNWHFLPH....DSWEETSAKIH
Zma IVPSISNFSSSYYLLTHNQILLKKYLFLDNLKQTLQVLQGLKYSLIDENKRISNFD..SNIMLEPFHLNWHFLHH....DSWEETLAIIH
SBHM8 SBHM2 TLc
920 930 940 950 960 970 980 990 1000
Sel PGSLIVAGTQLVDGIIAPESGEVRAIAP....GQLQLRIARPYRVSQGAVLHVEDKGLVQRGDNLVLLVFERAKTGDIIQGLPRIEELLE
Tel VGDLVRPGDELAKGVTAPETAAVMAVAD....DHVILRLARPYLVSPGAVLQIEEGDLVQRGDNLALLVFERAKTGDIIQGLPRIEELLE
Syn PGQLLVAGQELVPGVKLEESGKVLEINGKGDNYQLVLRRARPYRVSPGAVLHIEDGDLVQRGDNLVLLVFERAKTGDIVQGLPRIEELLE
Ath LGQFFCENVCIAKKEPHLKSGQVLIVQR...DSAVIRSAKPYLATPGAKVHGHYSEILYEGDTLVTFIYEKSRSGDITQGLPKVEQVLE
Osa LGQFICENVCLFKSH.IKKSGQIFIVNI....DSFVIRAAKPYLATTGATVHGHYGEILYKGDRLVTFIYEKARSSDITQGLPKVEQIFE
Zma LGQFICENLCLFKLH.IKKSGQIFIVNM....DSFVIRAAKPYLATIGATVHGHYGKILYKGDRLVTFIYEKSRSSDITQGLPKVEQIFE

**SFig. 3.** Si3 structure and sequence alignment. (A) The structure of elongation complex with NusG are shown as transparent surface, and the Si3, DNA/RNA and trigger loop ($TL_N$, $TL_C$) are shown as cartoon models. The SBHM are labeled 1 to 8 and sub-domains (Tail, Fin, Body and Head) are indicated. The active site of RNAP is indicated as it coordinating $Mg^{2+}$ (magenta sphere). (B) Sequence alignment comparing cyanobacterial Si3 (Sel, *Synechococcus elongatus*; Tel, *Thermosynechococcus elongatus*; Syn *Synechocystis* sp. PCC 6803) with the corresponding inserts from chloroplast RNAP of plant (Ath, *Arabidopsis thaliana*; Osa, *Oryza sativa*; Zma, *Zea mays*). The identical and conserved sequences are indicated by red background and red text, respectively. The sequence alignment was performed using Clustal-Omega (44) and visualized by Endscript 2.0 (45). The number scale above the sequences corresponds to the Sel sequence. Insertions found in chloroplast Si3 domains are indicated by Ω.

SFig. 4

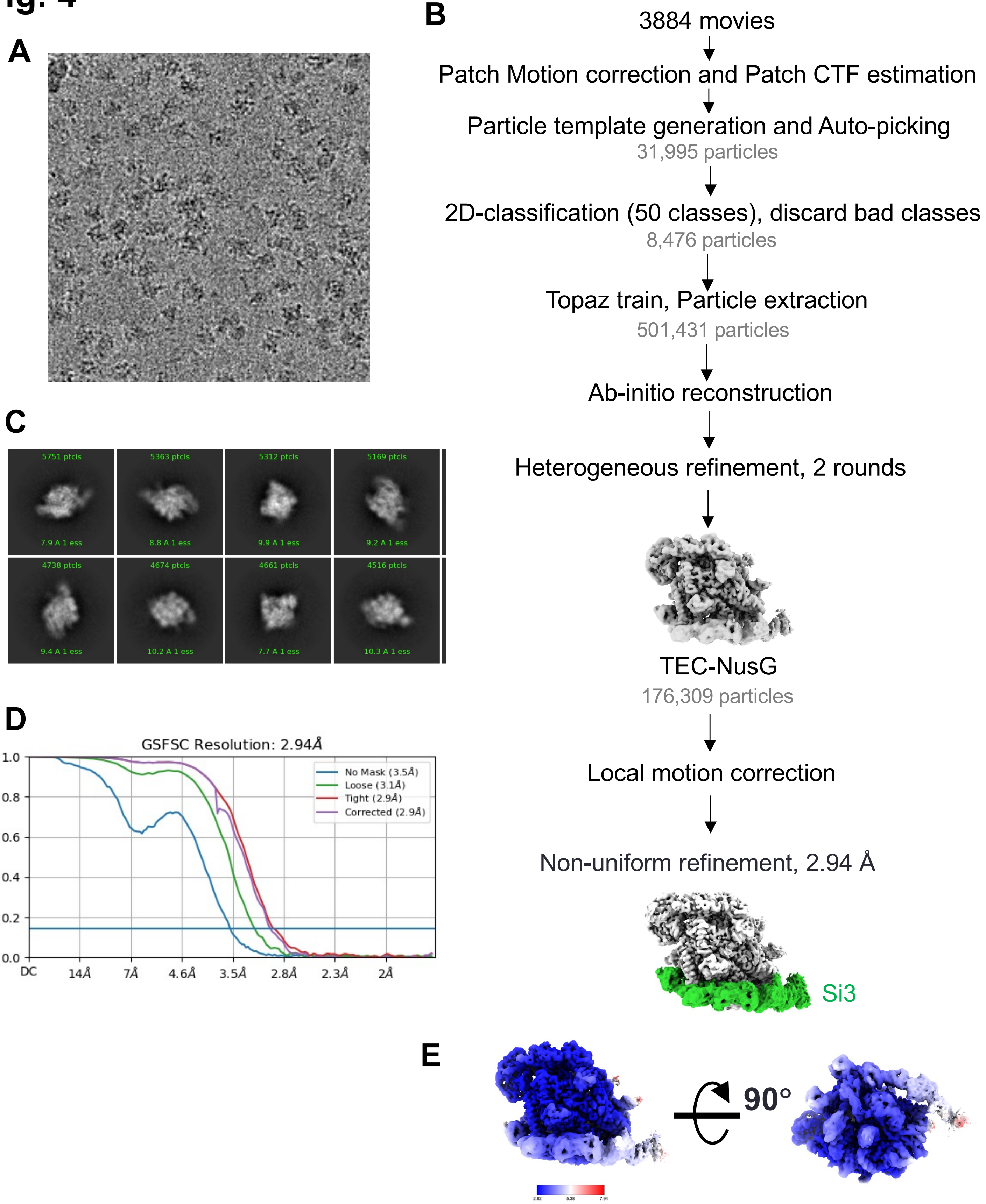


**SFig. 4. Cryo-EM processing pipeline of the cyRNAP elongation complex with NusG.** (A) A representative micrograph used for data processing. (B) Cryo-EM data processing and classification flow chart. (C) Selected representative 2D classes. (D) Fourier shell correlation (FSC) plot for half-maps with 0.143 FSC criteria indicated. (E) Local resolution maps.

### SFig. 5

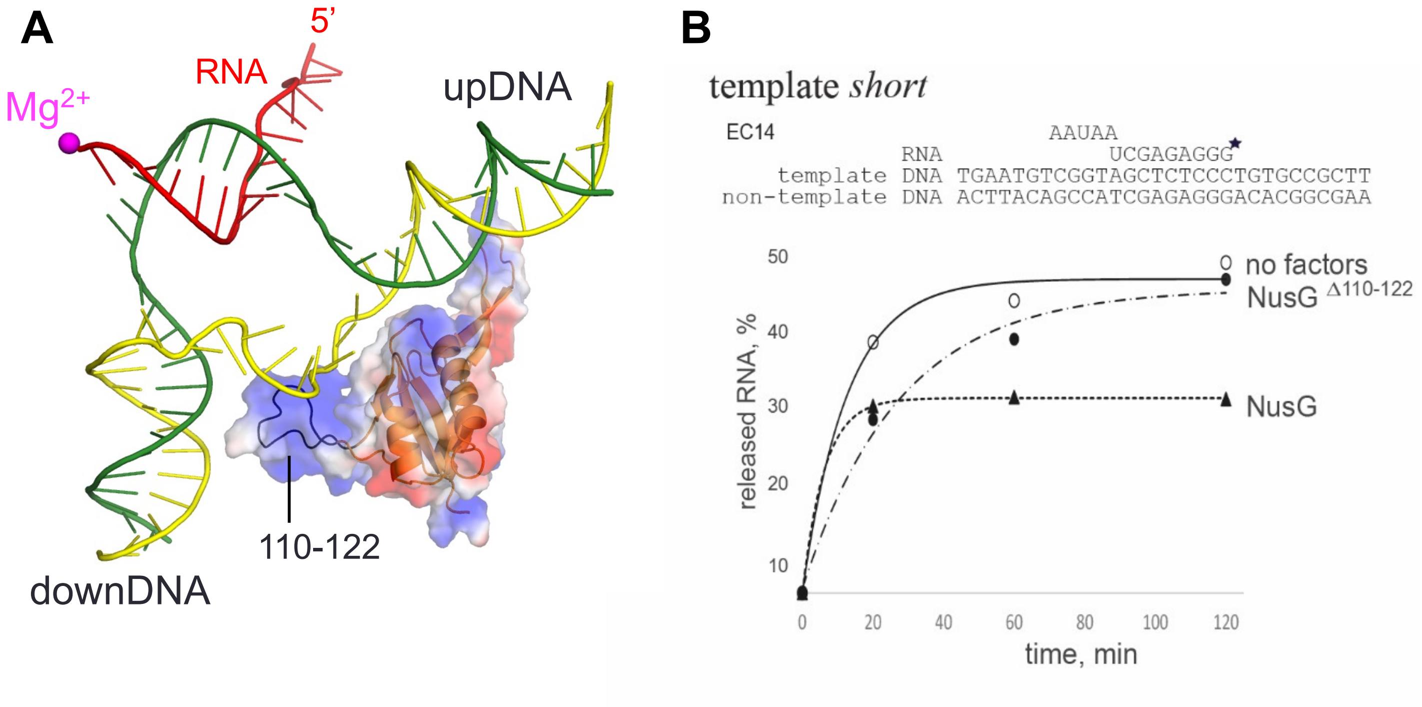


**SFig. 5. Structure and function of cyanobacterial NusG.** (A) NusG interaction with DNA. DNA/RNA scaffold and NusG (orange) from the EC-NusG are depicted with cartoon models with transparent surface of NusG showing its electrostatic potential (blue and red, positive and negative). A unique loop of cyanobacterial NusG (residues 110-112) is indicated by black. (B) NusG increases stability of cyRNAP elongation complex. Elongation complex was assembled using oligo DNA and RNA shown above the plot, and immobilized on Ni-NTA sepharose beads via His6 tag on RNAP. Plot shows kinetics of RNA release from the elongation complexes challenged with 300 mM NaCl, in percentage from total initial amount of RNA in complex.

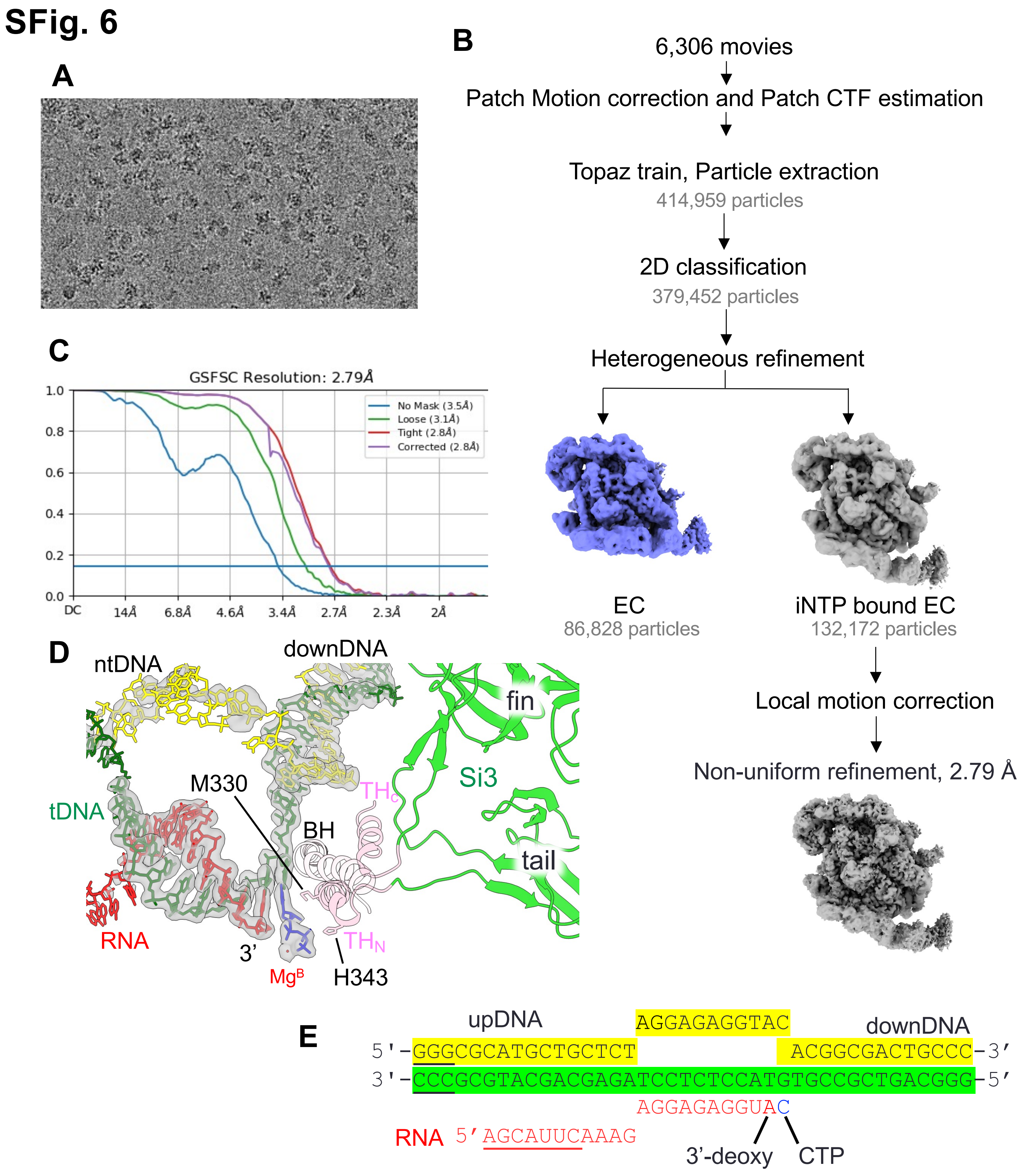
SFig. 6
A
B
6,306 movies
Patch Motion correction and Patch CTF estimation
Topaz train, Particle extraction
414,959 particles
2D classification
379,452 particles
Heterogeneous refinement
EC
86,828 particles
iNTP bound EC
132,172 particles
Local motion correction
Non-uniform refinement, 2.79 Å
C
GSFSC Resolution: 2.79Å
No Mask (3.5Å)
Loose (3.1Å)
Tight (2.8Å)
Corrected (2.8Å)
1.0
0.8
0.6
0.4
0.2
0.0
DC
14Å
6.8Å
4.6Å
3.4Å
2.7Å
2.3Å
2Å
D
ntDNA
downDNA
fin
Si3
tail
tDNA
M330
BH
THC
THN
RNA
3'
MgB
H343
E
upDNA
AGGAGAGGTAC
downDNA
5'-GGGCGCATGCTGCTCT ACGGCGACTGCCC-3'
3'-CCCGCGTACGACGAGATCCTCTCCATGTGCCGCTGACGGG-5'
AGGAGAGGUAC
RNA 5'AGCAUUCAAAG
3'-deoxy
CTP

**SFig. 6. Cryo-EM processing pipeline of the iNTP bound cyRNAP elongation complex with NusG.** (A) A representative micrograph used for data processing. (B) Cryo-EM data processing and classification flow chart. (C) Fourier shell correlation (FSC) plot for half-maps with 0.143 FSC criteria indicated. (D) The structure of active site of iNTP bound EC overlayed with cryo-EM map of DNA, RNA, iNTP and Mg (gray transparent). (E) The sequence of DNA/RNA scaffold used for preparing the iNTP bound EC-NusG (template DNA, green; non-template DNA, yellow; RNA, red). 3'-deoxy modified RNA was prepared by extending RNA with 3'-deoxy ATP after assembled EC with cyRNAP, DNA/RNA and NusG. DNA and RNA regions lacking cryo-EM density are underlined.

### SFig. 7

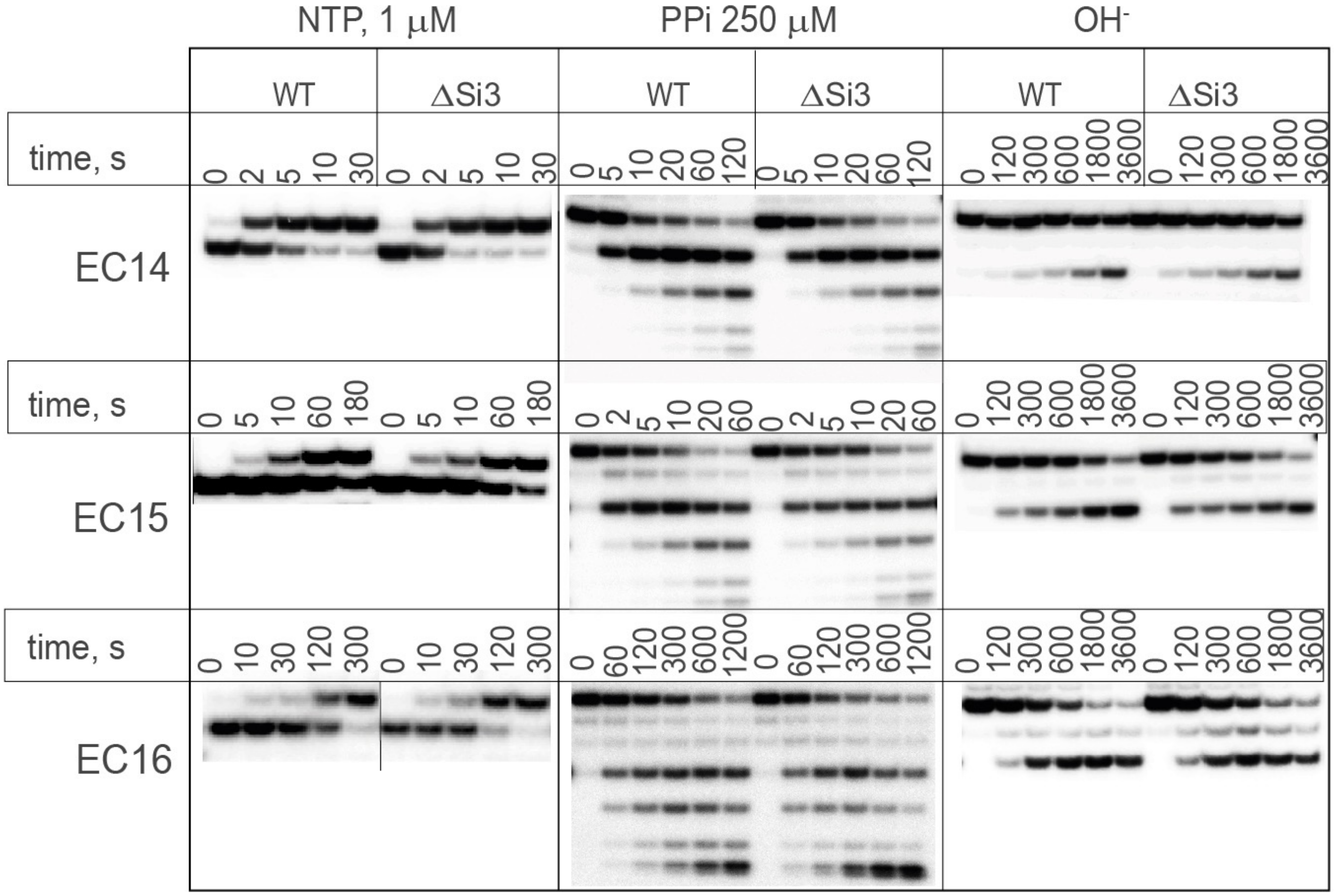


**SFig. 7. Activity of WT and ΔSi3 cyRNAPs.** Kinetics of single NTP incorporation, pyrophosphorolysis and phosphodiester bond hydrolysis (dinucleotide cleavage) in EC14, EC15 and EC16. Representative gels are shown. Related to Fig 4C.

**SMovie 1. Cryo-EM density map of the cyRNAP elongation complex with NusG.** Related to Fig. 2B.

**SMovie 2. Conformational changes of trigger loop/helix, rim helix and Si3 during the iNTP binding at the active site.** Related to Fig. 4B.
