## Supplementary material for "Structure and function of the Si3 insertion integrated into the trigger loop/helix of cyanobacterial RNA polymerase": SI Tables

**Supplemental Table 1.** X-ray crystallographic data collection and structure refinement statistics of the TelSi3ΔN

| **PDB code** | 8EMB |  |  |
| --- | --- | --- | --- |
| **Data collection** |  | **Refinement** |  |
| Space group | P3(2)21 | Resolution (Å) | 50–3.06 (3.10-3.06)* |
| Cell dimensions |  | *R*_work_ | 0.249 (0.387)* |
| *a* (Å) | 178.663 | *R*_free_ | 0.278 (0.371)* |
| *b* (Å) | 178.663 | No. of atoms | 21,086 |
| *c* (Å) | 281.038 | No. of waters | 0 |
| *α, β, γ* (°) | 90, 90, 120 | R.m.s deviations |  |
| Resolution (Å) | 50–3.07 (3.12-3.07)* | Bond length (Å) | 0.014 |
| Total reflections | 1,165,515 | Bond angles (°) | 1.557 |
| Unique reflections | 96,754 (4,161)* | Clashscore | 17.63 |
| Redundancy | 12.0 (9.5)* | Ramachandran favored, % | 91.97 |
| Completeness (%) | 98.9 (85.8)* | Ramachandran outliers, % | 0.07 |
| *I* / σ | 17.0 (0.74)* |  |  |
| CC^1/2^ | 0.999 (0.645)* |  |  |
| No. of Se sites | 44 |  |  |
| FOM | 0.753 |  |  |

*Highest resolution shells are shown in parentheses

**Supplementary Table 2. Cryo-EM data collection, refinement and validation statistics**

|  | **EC-NusG**  **(EMD-40874)**  **(8SYI)** | **EC-NusG_CTP**  **(EMD-** **42502)**  **(8URW)** |
| --- | --- | --- |
| **Data collection and processing** |  |  |
| Magnification | 75,000 | 105,000 |
| Voltage (kV) | 300 | 300 |
| Electron exposure (e**^-^**/Å^2^) | 45 | 40 |
| Defocus range (μm) | -0.75 to -2.5 | -0.75 to -1.75 |
| Pixel size (Å) | 0.87 | 0.855 |
| Symmetry imposed | C1 | C1 |
| Initial particle images (no.) | 501,431 | 414,959 |
| Final particle images (no.) | 176,309 | 132,172 |
| Map resolution (Å)  FSC threshold | 2.94  0.143 | 2.79  0.143 |
| Map resolution range (Å) | 2.82 -11.48 | 2.47 - 8.67 |
| **Refinement** |  |  |
| Initial model used (PDB code) | 8EMB, 8GZG |  |
| Model resolution (Å)  FSC threshold | 2.94  0.143 | 2.79  0.143 |
| Map sharpening *B* factor (Å^2^) | -109 | -53.4 |
| *Model composition*  Non-hydrogen atoms  Protein residues  Nucleic acid residues  Ligands | 29,500  3,537  91  Zn:2, Mg:1 | 29,896  3,574  94  Zn:2, Mg:1 |
| *B factors (Å^2^)* (mean)  Protein  DNA/RNA  Ligand | 71.71  171.97  100.75 | 106.22  195.73  61.08 |
| *R.m.s. deviations*  Bond lengths (Å)  Bond angles (°) | 0.005  0.644 | 0.003  0.562 |
| *Validation*  MolProbity score  Clash score  Rotamer outliers (%) | 2.46  6.85  6.09 | 2.26  6.27  4.41 |
| *Ramachandran plot*  Favored (%)  Allowed (%)  Disallowed (%) | 91.95  7.74  0.31 | 93.54  6.12  0.34 |
